# MS4A8B regulates Orai1-dependent Ca²⁺ influx to control motile cilia function in human nasal epithelial cells

**DOI:** 10.64898/2026.08.17.745248

**Authors:** Alexander A. Simon, Ray Z. Ma, Jerry S. Rao, Katherine Rozsypalek, Zhongming Ma, Nithin D. Adappa, James N. Palmer, Yobouet Ines Kouakou, Robert J. Lee

## Abstract

Motile cilia demonstrate coordinated beating to propel fluids across epithelial tissues, and changes to their beating frequency are largely regulated by intracellular second messengers including Ca^2+^. In the airway epithelium, ciliary beating is essential to mucociliary clearance. Mucociliary clearance involves trapping inhaled pathogens and irritants in sticky mucus lining the airways for motile cilia to sweep away contaminated mucus, preventing infection and reducing general airway inflammation. Many chronic respiratory diseases, including chronic rhinosinusitis and asthma, are characterized by an acquired ciliary dysfunction. Despite the importance of Ca^2+^ signaling in cilia physiology, the identity and molecular mechanisms governing localized ciliary Ca^2+^ transport remain poorly understood. MS4A8B is an uncharacterized cilia-localized transmembrane protein. Other MS4A homologs have been indirectly linked to Ca^2+^ signaling via uncharacterized mechanisms. Using primary human nasal epithelial cells differentiated at air-liquid interface, we demonstrated that MS4A8B regulates motile cilia function. MS4A8B knockdown impairs ciliary beating and impacts cilia structure. Live-cell imaging combined with genetic analysis revealed that MS4A8B potentiates Orai1-mediated Ca^2+^ influx. Co-immunoprecipitation and FRET microscopy in ectopic expression systems demonstrated that MS4A8B interacts with Orai1 channels. Orai1 was further identified to reside in motile cilia of primary human nasal epithelial cells, allowing ciliary beat frequency to be stimulated by Orai1 agonists including arachidonic acid. MS4A8B functional coupling with Orai1 acts as an autonomous cilia signaling network. Targeting this compartmentalized signaling pathway offers a novel therapeutic approach to restore or enhance mucociliary clearance in airway diseases.

**Highlights:**

- Loss of MS4A8B expression in airway motile cilia leads to ciliary dysfunction
- MS4A8B regulates Ca^2+^ signaling through potentiation of Orai1 Ca^2+^ influx
- Orai1 is a motile cilia localized Ca^2+^ channel
- Ciliary beat frequency and mucociliary clearance are stimulated by arachidonic acid

## Introduction

Motile cilia are specialized organelles acting as microscopic propellers that rhythmically beat to accomplish fluid’s mechanical movement. Motile cilia exist in the respiratory tract, reproductive tracts, and brain ventricles, where they extend from epithelial cell’s plasma membrane into the surrounding environment. In the airway epithelium, the clearance of mucus by cilia is a front-line defense against inhaled pathogens and irritants.^1^ Sticky mucus traps harmful foreign bodies, and motile cilia sweep away contaminated mucus.^2^ Ciliary dysfunction is a hallmark of many chronic respiratory diseases, including chronic rhinosinusitis, chronic obstructive pulmonary disease, and asthma.^3^ ^4^ Altered cilia function is also linked to susceptibility to acute infection or irritation by environmental pollutants.^1^ ^3^ In the reproductive tracts, motile cilia propel gametes and impact fertility,^5,6^ while brain motile cilia circulate cerebrospinal fluid.^7^

Ca^2+^ is a conserved key second messenger that modulates motile cilia beating in airways^8,9^, ependymal neurons^10^, and fallopian tubes^11,12^. Primary cilia, which are non-motile chemosensory antennae, can autonomously regulate their own ciliary Ca^2+^ gradients through ciliary membrane Ca^2+^ channels.^13,14^ Similarly, flagella from sperm—which are structurally analogous to motile cilia—also rely on Ca^2+^ permeable ion channels in various eukaryotic organisms.^15-17^ Although there is well-established evidence of Ca^2+^ regulating motile ciliary beat frequency (CBF), the molecular genetics demonstrating Ca^2+^ transport mechanisms in motile cilia is incomplete.^18-21^

We sought to investigate a member of the membrane-spanning four-domain subfamily A (MS4A) protein family identified as MS4A8B and evaluate its function in airway epithelial cilia signaling. MS4A8B expression increases with nasal epithelial cell differentiation^22^ and has been localized to motile cilia across the nasopharynx, bronchus, fallopian tube, endometrium, and cervix.^23,24^ However, no functional data regarding MS4A8B is currently available. Other MS4A homologs have been implicated in Ca^2+^ signaling through undefined mechanisms.^25-32^ Cryo-EM of human MS4A1, also known as CD-20, revealed an oligomeric dimer lacking a central ion channel pore.^33^ Mouse MS4A homologs may be putative chemoreceptors that activate ligand-dependent Ca^2+^ influx in olfactory neurons.^34,35^ While these reports propose new hypotheses about how MS4A homologs’ trigger Ca^2+^ signaling in a variety of cell types, their exact mechanisms remain unclear.

We hypothesized MS4A8B controls motile cilia function by regulating Ca^2+^ signaling. To test this rigorously, we used live cell imaging and electrophysiological techniques in both cell lines and primary human nasal epithelial cells differentiated at air-liquid interface to model human motile cilia. We identified a novel mechanism of MS4A8B regulation of Orai Ca^2+^ influx detailed below.

## Results

### MS4A8B’s ciliated cell expression and predicted ciliary function

Publicly available single-cell RNA sequencing from nasal epithelial cells show MS4A8B expression within upper airways is exclusive to ciliated cells (Figure 1A). Comparative analysis of MS4A8B and the other 17 MS4A homologs further demonstrates MS4A8B’s ciliated cell expression is a unique characteristic among the MS4A family. No other MS4A homolog is detected in nasal ciliated cells, but most predominantly demonstrate myeloid expression patterns (Figure 1B).^36-38^ Cross-examination of single-cell gene expression from separate analyses of nasopharynx and lung validated MS4A8B’s ciliated cell exclusivity (Figure S1A,B).

**Figure 1:**
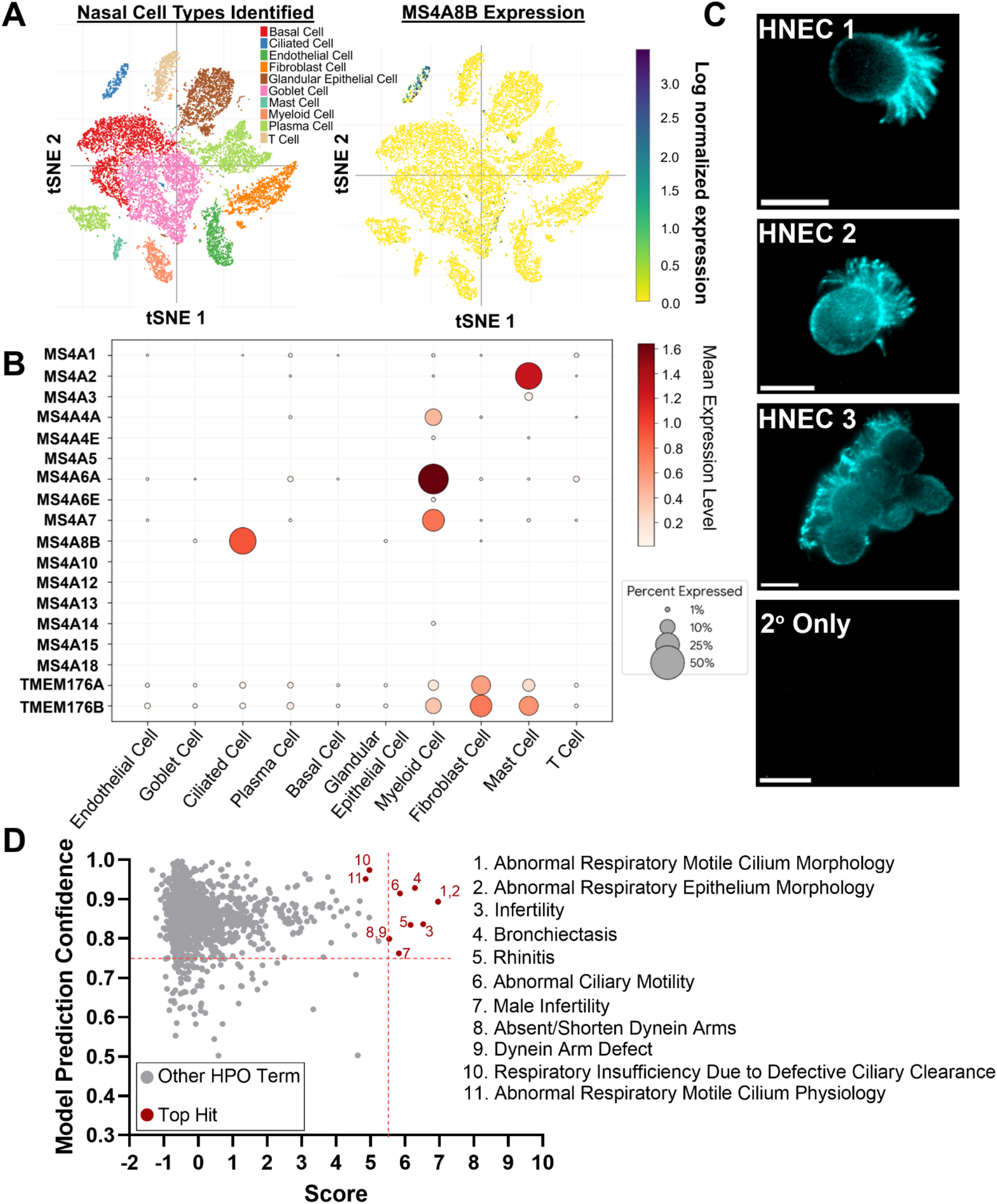
Ciliated cell-specific expression of MS4A8B in human nasal epithelial cells and predicted role in ciliary function. (A) Two-dimensional tSNE projections (18,036 cells) depicting distinct cell clusters (left) and corresponding MS4A8B expression (right). Data extracted from Broad Institute’s Single Cell Portal.^78,79^ (B) Dot plot comparing gene expression of all MS4A homologs in human nasal epithelial cells. (C) Immunofluorescence of isolated primary HNECs from three patients labeled for MS4A8B, with representative secondary-only control. Scale bar = 10μm. (D) Top enriched Human Phenotype Ontology (HPO) terms corresponding to MS4A8B from ARCHS4 database co-expression predictions.^39,80^ Top hits selected by upper quartile prediction confidence and score (providing a metric for statistical significance) >5.5. See also Figures S1, S2, and S3.

We therefore tested MS4A8B’s subcellular localization in primary human nasal epithelial cells (HNECs) from differentiated air-liquid interface (ALI) cultures grown from residual patient surgical material. Immunofluorescence revealed MS4A8B protein preferentially localized to motile cilia relative to basolateral membranes of isolated HNECs (Figure 1C), and intact HNEC monolayers (Figures S2 and S3).

To predict functional roles and clinical phenotypes associated with MS4A8B, gene-gene co-expression matrices were analyzed in PrismEXP.^39^ The top-scoring functional predictions heavily converged on airway clearance mechanisms and motile cilia structural and function (Figure 1D). These data suggest MS4A8B is a specific marker for ciliated cells in the upper airway and may play a critical role in maintaining motile ciliary function and mucociliary clearance.

### MS4A8B knockdown results in impaired motile cilia

We tested the effect of MS4A8B gene knockdown in primary HNEC ALIs using antisense oligonucleotides (ASOs).^40^ ASOs targeting MS4A8B (*ms4a8b*) delivered to polarized ALIs reduced both mRNA (Figure 2A) and protein (Figure 2B) expression. Concomitantly, we observed a reduction in the percent area of beating cilia from unstimulated ALI cultures treated with *ms4a8b* compared to scramble controls (Figure 2C). Cilia beating faster than 2.0 Hz was considered actively beating, whereas regions below this threshold were deemed inactive. Upon ATP stimulation, *ms4a8b* cultures showed a marginal response in the recruitment of beating cilia compared to scramble controls (Figure 2D,E). However, ciliary beat frequency (CBF) increases were similar (Figure 2F). Application of extracellular ATP mobilizes Ca^2+^ from both intracellular stores downstream of P2Y receptors and stimulates Ca^2+^ influx via P2X receptors.^41^ The lack of difference in maximal CBF after ATP suggests any functional motile cilia after ASO transfections maintained uninterrupted purinergic receptor pathways. The difference in the recruitment of beating cilia after purinergic stimulation suggests MS4A8B knockdown was associated with either the formation of some cilia unable to beat or an overall motile cilia degradation. To test the integrity of motile cilia differentiation, we performed immunofluorescence of polarized HNECs to examine microtubule presence. Scramble-treated ALIs showed abundant cilia staining reflected by β-tubulin IV (β-Tub IV), while MS4A8B knockdown revealed regions where β-Tub IV was not detected, indicating intact cilia were degraded (Figure 2G,H). These findings suggest MS4A8B-dependent signaling maintains motile cilia function, thus promoting efficient mucociliary clearance.

**Figure 2:**
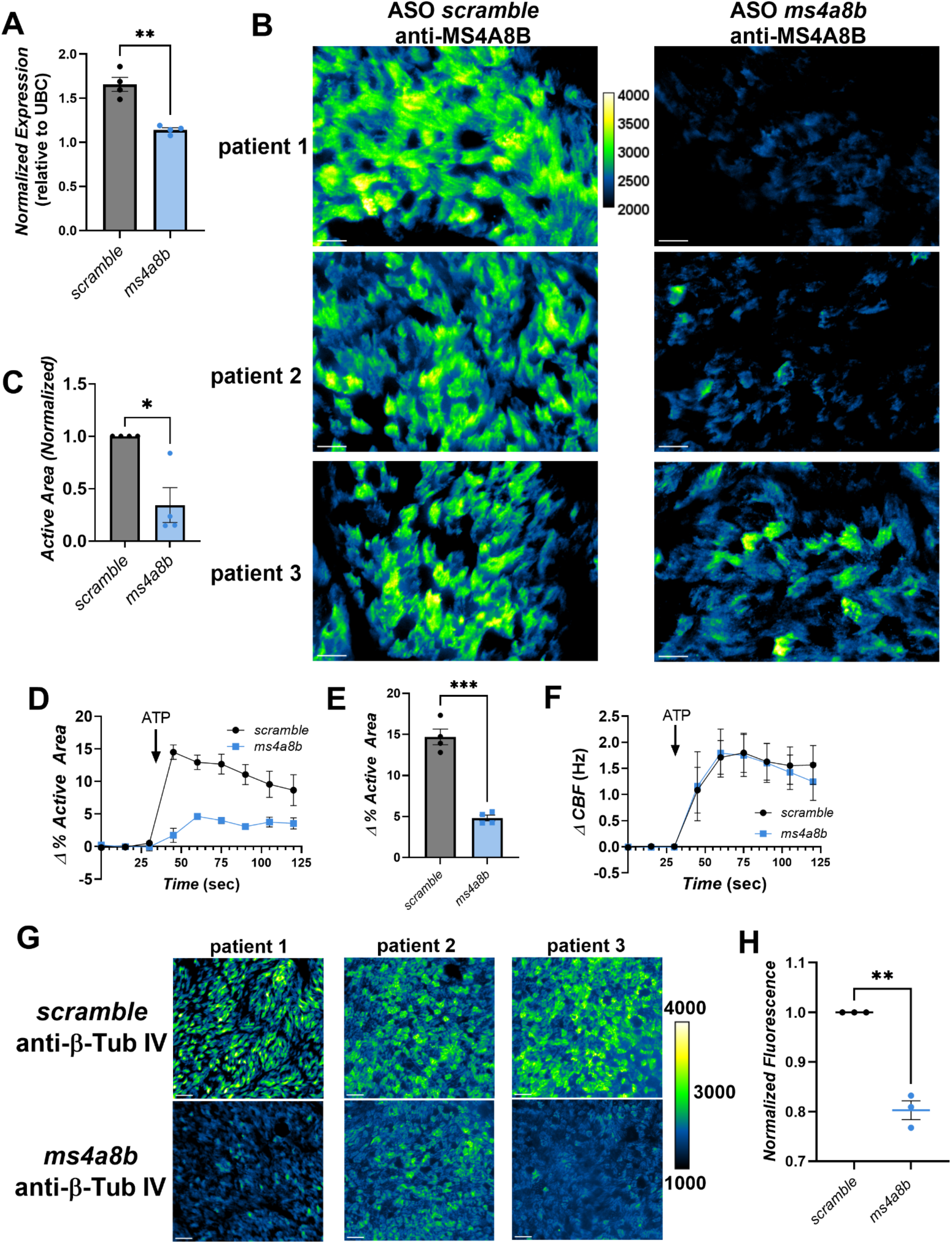
MS4A8B knockdown in primary HNECs impairs cilia structure and basal activity. (A) qPCR for MS4A8B in primary HNECs transfected with scramble or MS4A8B targeting antisense oligonucleotides (ASOs) (mean ± SEM). n=4 independent patients. Significance by Welch’s unpaired t-test, **p=0.0049. (B) Z-projections from primary HNECs labeling MS4A8B in ASO transfected primary HNECs. n=3 independent patients. Scale bars=20μm. (C) Ciliary beating measurements from primary HNECs showing percent area of ciliary beating under resting conditions (mean ± SEM, n=4 independent patients). Significance by Welch’s unpaired t-test, *p=0.0288 (D) Time course of change in area of beating cilia (mean ± SEM) after ATP (100μM) between scramble ASO and MS4A8B targeting ASO. n=4 independent patients. (E) Summary statistics (mean ± SEM) from experiments in (D) showing individual changes in percent area of beating cilia evaluated by paired t-test, ***p=0.0007. (F) Time course of change in CBF (mean ± SEM) after ATP (100μM) between scramble ASO and MS4A8B targeting ASO. n=4 independent patients. (G) Z-projections of primary HNECs comparing β-Tub IV in scramble and MS4A8B targeting ASO. n=3 independent patients. Scale bars=40μm. (H) Quantification of fluorescence intensity (mean ± SEM) corresponding to images in (G) evaluated with Welch’s unpaired t-test, **p=0.0092.

### MS4A8B functionally regulates ER Ca^2+^ store capacity

To characterize MS4A8B’s signaling mechanism in the airway epithelium, we took advantage of MS4A8B’s expression in pulmonary neuroendocrine cells (PNECs), a rare lung cell type that is also the cell type of origin for small cell lung cancer (SCLC) (Figure S4). We used three independent PNEC-derived SCLC lines (DMS53, DMS454, and DMS153) that endogenously express MS4A8B. In all three lines, siRNA targeting MS4A8B (si*MS4A8B*) reduced MS4A8B mRNA expression (Figure S5). We then challenged SCLC cells to carbachol (CCH) to assess MS4A8B’s participation in Ca^2+^ signaling.^42^ CCH activated Ca^2+^ signals in all three cell lines when transfected with non-coding siRNA (NC-1), but all responses were attenuated by si*MS4A8B* transfection (Figure 3A,B,C, and Figures S6A,B, S6E,F). To challenge total ER Ca^2+^ stores, we applied SR/ER Ca^2+^-ATPase (SERCA) inhibitor thapsigargin (TG) in the absence of extracellular Ca^2+^ to visualize ER Ca^2+^ leak, which informs on ER Ca^2+^ content. We observed attenuated TG-induced Ca^2+^ signals with MS4A8B knockdown. This suggests MS4A8B loss resulted in depleted ER Ca^2+^ stores (Figures 3D,E, S6C,D, S6G,H). Supporting this, ER Ca^2+^ concentration ([Ca^2+^]_ER_) measured by the CaMeleon-based probe D1ER^43^ indicated lower [Ca^2+^]_ER_ in si*MS4A8B*-treated cells (Figure 3F).

**Figure 3:**
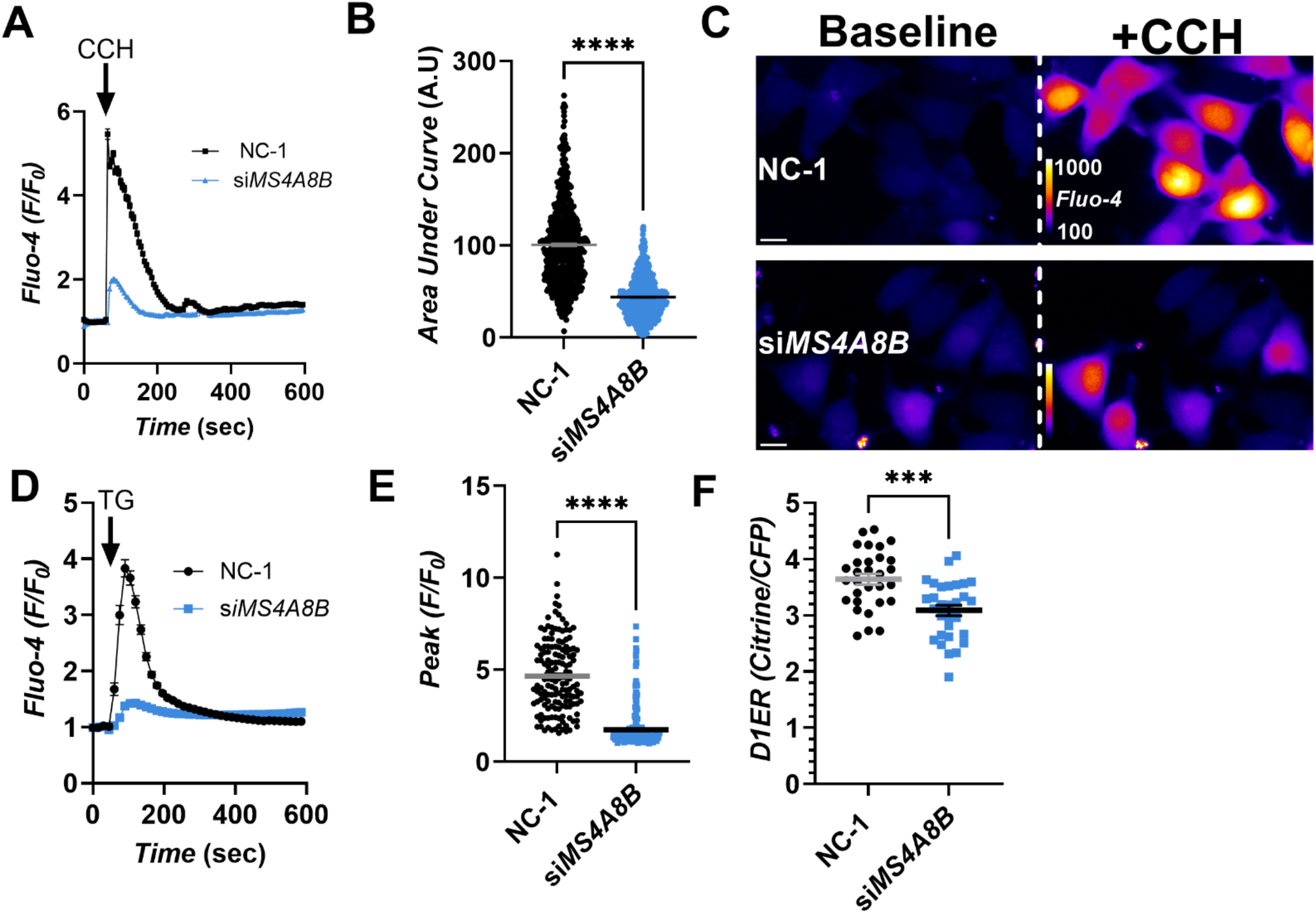
MS4A8B regulates ER Ca^2+^ stores in airway epithelial cells. (A) [Ca^2+^]_cyt_ time course reported by Fluo-4 (mean ± SEM) in DMS53 cells transfected with non-coding siRNA (NC-1, n=625 cells) or si*MS4A8B* (n=608 cells) treated with carbachol (CCH) (10μM). (B) Summary statistics (mean ± SEM) from (A) showing area under the curve (AUC), defined in arbitrary units (A.U), from individual cell’s [Ca^2+^]_cyt_ traces. NC-1, n=622; si*MS4A8B* n=597, outliers were identified by ROUT (Q=1%). Significance by Welch’s unpaired t-test, ****p=<0.0001. (C) Representative pseudo-colored images depicting Ca^2+^ responses from (A). Scale bar indicates 10μm. (D) [Ca^2+^]_cyt_ time course by Fluo-4 (mean ± SEM) in DMS53 cells transfected with NC-1 (n=149 cells) or si*MS4A8B* (n=224 cells) treated with thapsigargin (TG) (10μg/mL). Cells were imaged in Ca^2+^-free bath solution (1mM EGTA). (E) Summary statistics of (D) showing the peak Fluo-4 (mean ± SEM) of individual cells. Significance by Welch’s unpaired t-test, ****p=<0.0001. (F) D1ER reported ER Ca^2+^ content (mean ± SEM) in DMS53 cells between NC-1 (n=30 cells) and si*MS4A8B* (n=30 cells) evaluated with unpaired t-test, ***p=0.0001. See also Figures S4, S5, and S6.

### MS4A8B potentiates store-operated Ca^2+^ entry

While multiple mechanisms could explain the altered Ca^2+^ signaling signatures observed with MS4A8B knockdown—as MS4A8B is a plasma membrane protein—we hypothesized depleted [Ca^2+^]_ER_ may be due to the lost MS4A8B-dependent regulation of store-operated Ca^2+^ entry (SOCE).^25,27^ SOCE is required to replenish [Ca^2+^]_ER_ stores after ER Ca^2+^ release, and occurs when ER-resident protein STIM senses the fall of [Ca^2+^]_ER_ and activates plasma membrane Orai Ca^2+^channels to allow Ca^2+^ influx and refilling of ER Ca^2+^ stores by SERCA. When mCherry-MS4A8B was expressed in lung BEAS-2B cells, MS4A8B expression was observed on the plasma membrane (Figure S7). After TG-induced [Ca^2+^]_ER_ depletion, CaMeleon-based cytosolic Ca^2+^ indictor D3cpv reported greater Ca^2+^ influx after reintroduction of Ca^2+^ in MS4A8B expressing cells relative to cells transfected with cDNA coding for only free mCherry (Figure 4A,B,C).

**Figure 4:**
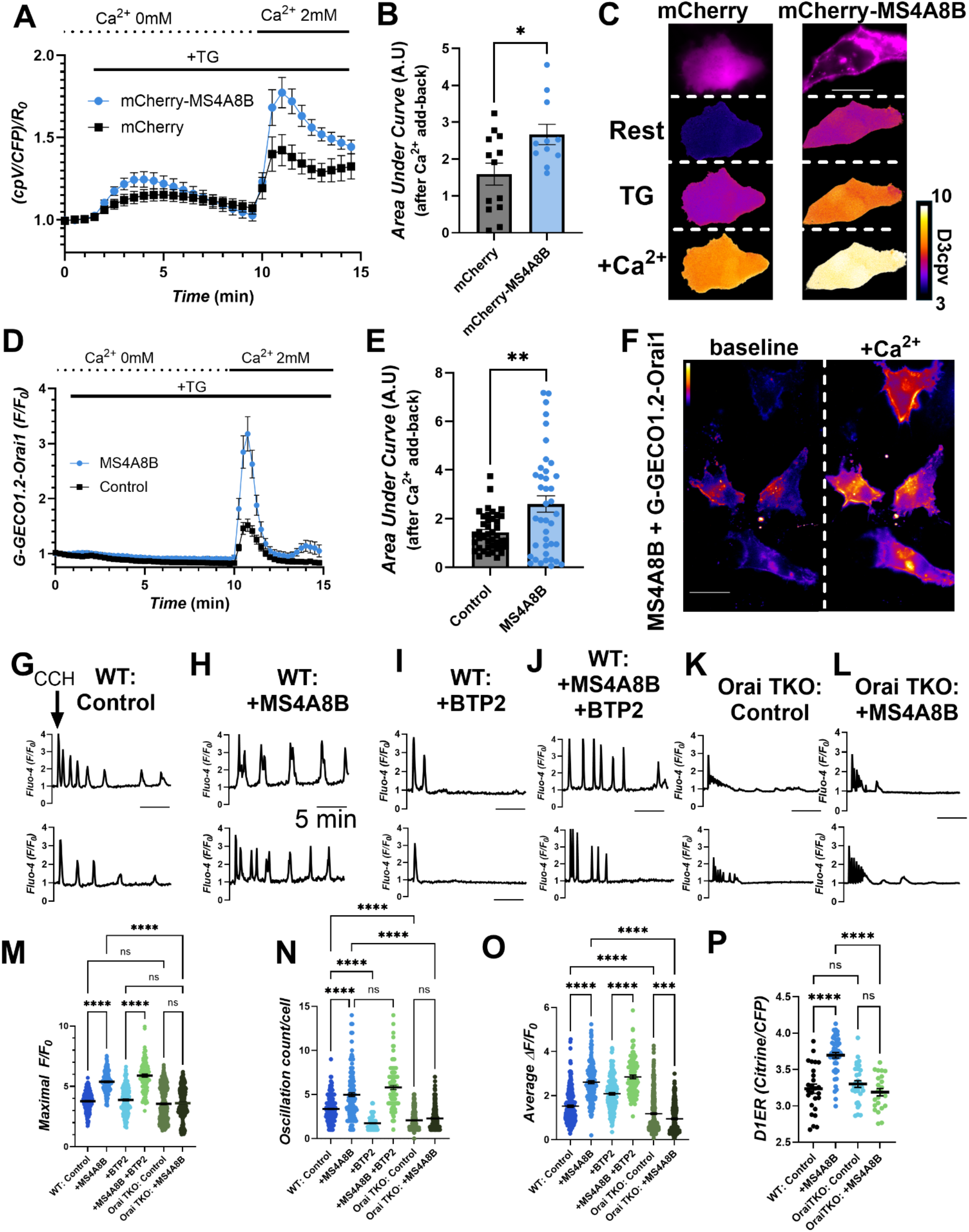
MS4A8B potentiates store-operated Ca^2+^ entry. (A) [Ca^2+^]_cyt_ time course reported by D3cpv (mean ± SEM) in BEAS-2Bs expressing free mCherry (n=13 cells) or mCherry-MS4A8B (n=11 cells) following ER Ca^2+^ store depletion with thapsigargin (TG) (10μg/mL) and Ca^2+^ add-back (2mM). (B) Summary statistics (mean ± SEM) from experiments in (A) reporting cell’s AUC, tested with Welch’s unpaired t-test, *p= <0.0152. (C) Representative images of [Ca^2+^]_cyt_ (D3cpv) from experiments in (A) at baseline, peak TG (4min), and peak add-back (11min). Scale bar indicates 25μM. (D) [Ca^2+^]_cyt_ time course (mean ± SEM) reported by G-GECO1.2-Orai1 in BEAS-2Bs co-transfected with MS4A8B-IRES-mCherry (MS4A8B, n=40 cells) or mCherry alone (Control, n=42 cells). (E) Summary statistics (mean ± SEM) of AUC from each cell in (D). Significance by Welch’s unpaired t-test, **p= <0.0021. (F) Representative images of G-GECO-Orai1 Ca^2+^ signals in BEAS-2Bs co-expressing MS4A8B at rest and following Ca^2+^ add-back. Scale bar indicates 25μM. (G-H) Representative Ca^2+^ oscillations (Fluo-4) induced by carbachol (CCH, 10μM) in wild-type (WT) HEK293s with (H) or without (G) MS4A8B expression. (I-J) Representative Ca^2+^ oscillations induced by CCH with BTP2 (10μM) in WT HEK293s with (J) or without (I) MS4A8B. (K-L) Representative CCH-induced Ca^2+^ oscillations in Orai triple knockout (TKO) HEK293s with (L), or without (K) MS4A8B. (M-O) Summary statistics (mean ± SEM) analyzing data from (G-L). Individual cells were analyzed for outliers by the ROUT test (Q=1%) and averaged over 3-5 independent passages. Significance by Brown-Forsythe and Welch ANOVA with Dunnett’s multiple comparison test. ****p=<0.0001, ***p=0.0002. (P). ER Ca^2+^ content (D1ER, mean ± SEM) analyzed using Brown-Forsythe and Welch ANOVA with Dunnett’s multiple comparison test ***p=<0.0001. WT control (n=26 cells), WT +MS4A8B (n=46 cells), Orai TKO control (n=31 cells), Orai TKO + MS4A8B (n=23 cells). See also Figures S7, S8, S9, and S10.

To confirm the role of Orai1 in MS4A8B-potentiated Ca^2+^ addback, we utilized the G-GECO1.2 Ca^2+^ indicator N-terminally fused to Orai1 to selectively measure Ca^2+^ influx through the fused Orai1 channel.^44^ Ca^2+^ addback responses were again increased in amplitude when BEAS-2B cells were co-transfected with MS4A8B-IRES-mCherry (Figure 4D,E,F). While ER store Ca^2+^ release with TG was visualized by our cytosolic Ca^2+^ probe D3cpv (Figure 4A), note this initial release was not visualized with G-GECO1.2-Orai1 (Figure 4D). G-GECO1.2-Orai1 only reports the Ca^2+^ signals occurring locally through Orai1, and suggests MS4A8B is potentiating Orai1 specific Ca^2+^ influx.

Classical SOCE assays inducing [Ca^2+^]_ER_ depletion by TG and stimulating cells with the re-introduction of Ca^2+^ generates maximal SOCE function. We evaluated MS4A8B’s SOCE modulation by monitoring more physiological Ca^2+^ signals induced by CCH in HEK293 cells. CCH activation of mAChRs generates inositol 1,4,5-trisphosphate (IP_3_), which activates IP_3_ receptors (IP_3_Rs) to provide the physiological stimulus to release Ca^2+^ and lower [Ca^2+^]_ER_, activating SOCE. Orai channels are known to play a critical role in controlling regenerative Ca^2+^ oscillations in HEK293s.^45^ MS4A8B expression in HEK293s by transient transfection potentiated CCH-induced Ca^2+^ signals relative to control (empty vector) cells (Figure 4G,H). Maximal CCH-induced Ca^2+^ signals were larger with MS4A8B, and MS4A8B-expressing cells became more oscillatory, with a larger mean amplitude of each oscillation (Figure 4M,N,O).

We next challenged the MS4A8B-potentiated CCH-induced Ca^2+^ oscillations with co-treatment of SOCE inhibitor BTP2 (Figure 4I,J). In wild-type (WT) HEK293s, maximal CCH-induced signals were unaffected by BTP2, but Ca^2+^ oscillations were almost completely abolished (Figure 4I). BTP2-treated HEK293s expressing MS4A8B showed greater oscillatory capacity compared to WT but not significantly different compared to MS4A8B-expressing cells without BTP2 (Figure 4J; Figure 4M,N,O). We hypothesized BTP2’s reduced effectiveness against MS4A8B over-expression was due to expanded [Ca^2+^]_ER_ yielding greater IP_3_R Ca^2+^ efflux. To test this hypothesis, we used Orai1,2,3 triple knockout (TKO) HEK293s^46^, in which CRISPR was used to delete all three Orai homologs. Here, we tested for differences in CCH-activated Ca^2+^ oscillations when expressing MS4A8B (Figure 4K,L). All CCH signaling metrics were diminished in Orai TKO cells (Figure 4M,N,O). Moreover, MS4A8B over-expression failed to rescue Ca^2+^ phenotypes from the loss of Orai channels, confirming Orai channels are required for MS4A8B-potentiated Ca^2+^ signaling.

We then measured [Ca^2+^]_ER_ by ER-localized Ca^2+^ indicator D1ER. As expected, D1ER signals were increased with MS4A8B expression in WT HEK293s, and this observation was neutralized in OraiTKO background (Figure 4P). Expanded ER Ca^2+^ stores in HEK293s expressing MS4A8B coincided with increased Ca^2+^ efflux from the ER to the cytosol stimulated by TG (Figure S8). We further challenged the requirement of Orai1 for MS4A8B-related phenotypes by testing maximal SOCE by TG using HEK cells. Control HEK293s transfected with MS4A8B showed similar potentiation of SOCE from the endogenous Orai channels as observed in BEAS-2Bs (Figure S9A,B) However, no SOCE was detected with MS4A8B expressing OraiTKO cells (Figure S9C,D).

Unlike previous studies which hypothesized MS4A isoforms could be ion channels ^26,29,47^, our data demonstrate MS4A8B is not an ion channel itself or at least is not sufficient to mediate Ca^2+^ conductance in the absence of an Orai homolog. To more definitively test this, we complemented our imaging data with patch clamp electrophysiology to measure plasma membrane ion currents. Whole-cell patch clamp recordings in Orai TKO cells supported the finding that MS4A8B alone cannot provide Ca^2+^ current (Figure S10). MS4A8B and Orai1 co-expression in Orai TKO cells revealed larger current amplitudes sensitive to Gd^3+^ over Orai1 alone (Figure S10). Collectively, these findings demonstrate MS4A8B regulates membrane transport with Orai1 as an ion channel. Orai1 is both necessary and sufficient for MS4A8B’s ability to enhance Ca^2+^ signaling.

### Orai1 and MS4A8B demonstrate protein-protein interaction

Due to the long-term modulation of resting ER Ca^2+^ stores by MS4A8B over-expression or knockdown, we hypothesized a direct interaction between MS4A8B and Orai1 rather than through intermediate signaling messengers. MS4A8B-CFP and Orai1-YFP were co-expressed as Förster resonance energy transfer (FRET) pairs. Both proteins colocalized to the plasma membrane of BEAS-2B cells (Figure 5A,B). YFP (acceptor) photobleaching increased CFP (donor) fluorescence, suggesting FRET was occurring under basal conditions and indicating these proteins reside in close proximity to one another (Figure 5C-G). FRET efficiency was calculated at 0.19±0.04, and the mean distance between CFP and YFP was estimated at 6.24 nm with R_o_=4.9.^48^

**Figure 5:**
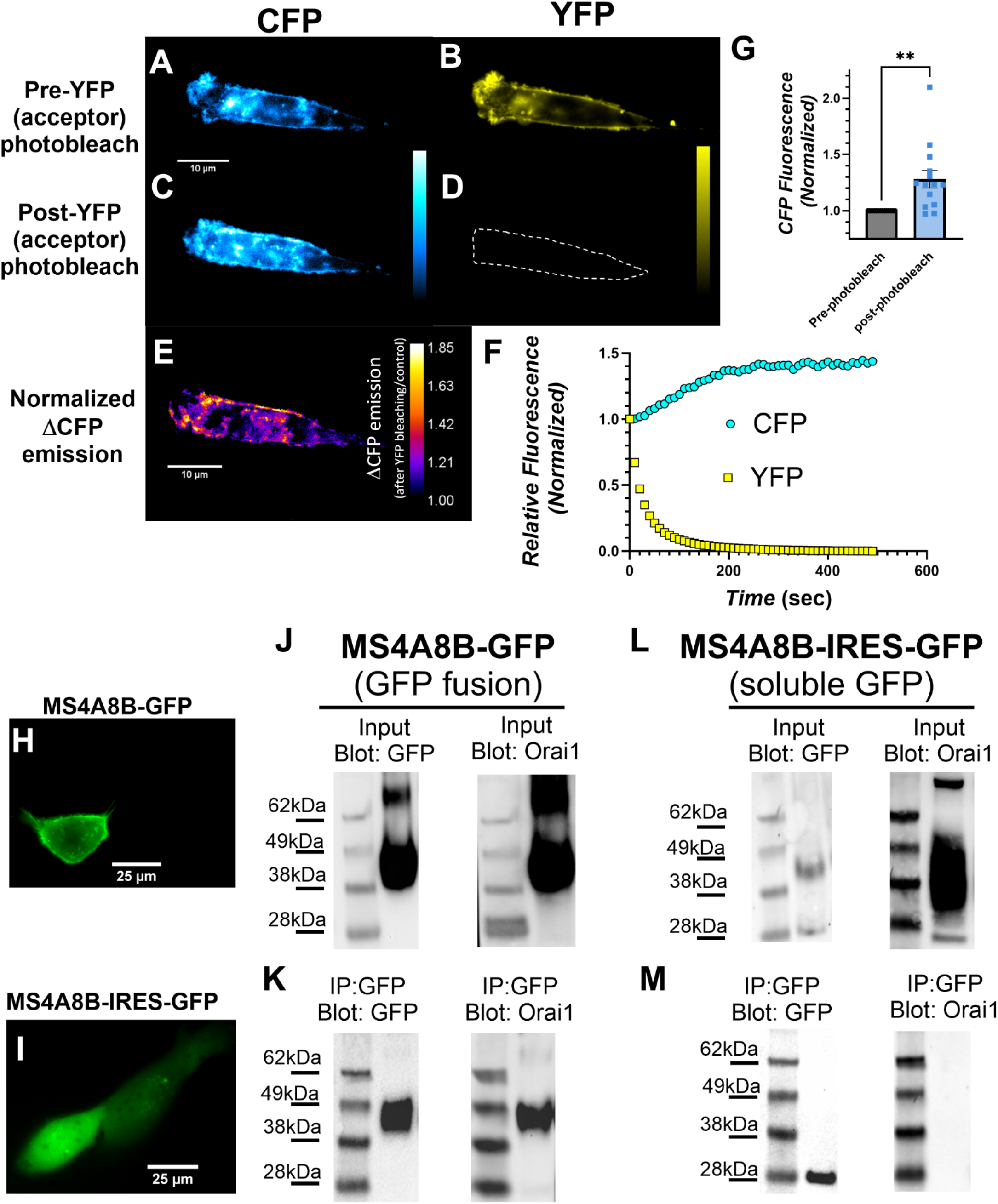
MS4A8B and Orai1 exhibit protein-protein interactions on plasma membrane. (A–D) Fluorescence images of BEAS-2Bs showing MS4A8B-CFP (donor) and Orai1-YFP (acceptor) intensity before (A, B) and after photobleaching (C, D) of the YFP acceptor. Dashed outline in D indicates photobleached region. E) Pseudocolor image of normalized increase in donor emission (ΔCFP emission) following YFP acceptor photobleaching relative to control. (F) Time course of relative fluorescence intensity (normalized) for CFP (cyan circles) and YFP (yellow squares) during photobleaching. (G) Quantification (mean ± SEM) of normalized CFP fluorescence intensity pre- and post-acceptor photobleaching tested with Welch’s unpaired t-test, **p=0.0035. (H-I) Localization of MS4A8B-GFP and MS4A8B-IRES-GFP in BEAS-2B cells. (J and K) Input control Western blots of GFP fusion constructs and endogenously expressed Orai1 from total BEAS-2B lysate. (L and M) Western blot for GFP or Orai1 from BEAS-2B lysate containing MS4A8B-GFP (L) or MS4A8B-IRES-GFP (M) after co-immunoprecipitation with selective GFP nanobodies.

We further tested our direct protein-protein interaction hypothesis using co-immunoprecipitation of Orai1 with a MS4A8B-GFP fusion construct and MS4A8B-IRES-GFP construct encoding a soluble GFP downstream of untagged MS4A8B. These constructs were transfected into BEAS-2B cells (Figure 5H,I). GFP was selectively pulled down by GFP nanobodies. Western blotting of protein extracted from MS4A8B-GFP-expressing cells revealed a band larger than expected for GFP alone and appropriately sized for MS4A8B fusion. In parallel, endogenous Orai1 was also detected by Orai1-antibody (Figure 5J,K). In MS4A8B-IRES-GFP-expressing cells, blotting for GFP revealed the expected 28kDa band for free GFP was a smaller molecular weight band than found with MS4A8B-GFP fusion, demonstrating a clean pull down free of non-specific co-immunoprecipitation. No Orai1 band was observed after GFP-pull down of lysate from MS4A8B-IRES-GFP cells (Figure 5L,M). This demonstrates the Orai1 protein found in the input control was lost by the selective GFP pulldown. Thus, MS4A8B-GFP was necessary for the pulldown of Orai1 with the GFP nanobodies.

### Orai1 localizes to motile cilia, and its activation can modify ciliary beating

The functionality of Orai Ca^2+^ channels has been demonstrated in SOCE in airway epithelial cells, but these studies are limited to non-differentiated, non-ciliated cells, including cell lines and primary cells grown in submersion rather than at air-liquid interface.^49,50^ Other studies suggest Orai1 expression is elevated in airway epithelial cells in inflammatory disease states, including asthma and cystic fibrosis, but with limited accompanying functional studies.^51^ Immunofluorescence of isolated primary HNECs to probe for the presence of Orai1 revealed labeling of basolateral regions and cilia (Figure 6A,B). This is expected as Orai channels’ ubiquitous role in SOCE likely necessitates expression of Orai1 in both apical and basolateral membranes, given Ca^2+^ signals can be distinctly compartmentalized in airway epithelial cells.^52^ Prior to the identification of Orai1 and STIM1, SOCE was observed in polarized ALI cultures evidence by [Ca^2+^]_cyt_ elevations in both apical and basolateral compartments after Ca^2+^ addback following ER Ca^2+^ depletion.^53^

**Figure 6:**
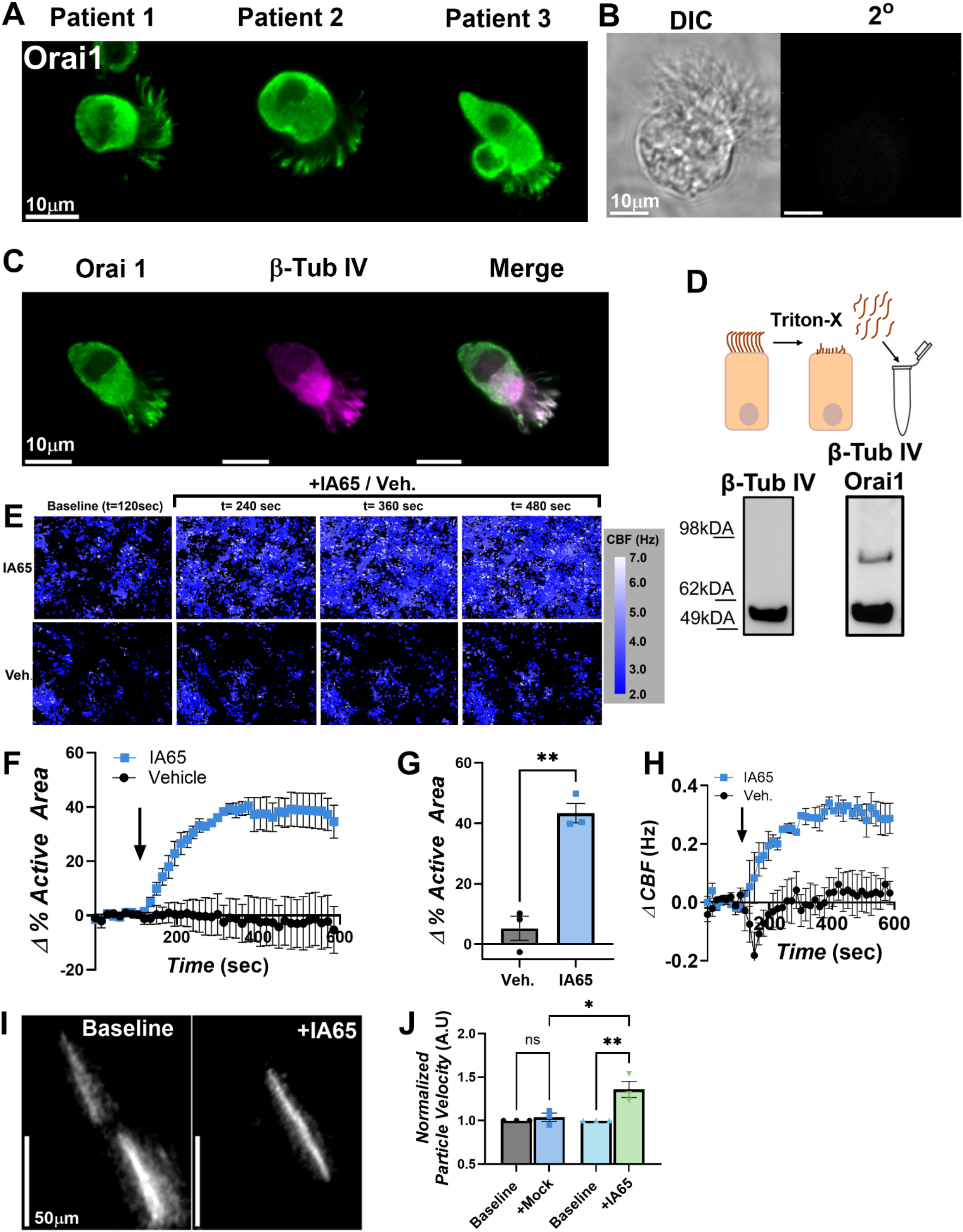
Orai1 is a motile cilia Ca^2+^ channel. (A) Orai1 localization by immunofluorescence in isolated primary HNECs from 3 independent patients. (B) DIC image and secondary antibody only control labeling of isolated primary HNECs. (C) Orai1 co-localization with β-Tub IV from primary HNEC. (D) Cartoon illustrating cilia harvesting and Western blot of pooled cilia membranes for β-Tub IV and Orai1. (E) Ciliary beating measurements with IA65 (1μM) treatment or mock vehicle treatment (0.05% DMSO) in primary HNECs illustrating active area of beating cilia and CBF of discrete regions. (F) Time course of change in area of beating cilia (mean ± SEM) after IA65 (1μM) or mock vehicle treatment (0.05% DMSO). n=3 independent patients. (G) Summary statistics of change on beating cilia area (mean ± SEM). Significance tested by paired t-test, **p=0.0065. (H) Time course of change in CBF (mean ± SEM) after IA65 (1μM) or mock vehicle treatment (0.05% DMSO). n=3 independent patients. (I) Particle transport micrographs from HNEC ALI cultures before and after IA65 (1μM). (J) Summary statistics of particle transport assay (mean ± SEM). For each condition, n=3 independent patients. Significance tested with a one-way ANOVA with Tukey’s multiple comparisons test, **p=0.0053, *p=0.0104. See also Figure S11.

Co-staining with motile cilia marker β-Tub IV showed overlapping patterns of expression with Orai1, suggesting at least some Orai1 is present in the ciliary membrane (Figure 6C). We validated Orai1 expression in cilia by Western blot of cilia specific membranes harvested from HNECs using an established protocol.^54^ Cilia membrane samples were enriched with β-Tub IV. Orai1 was also identified at the molecular weight consistent with a glycosylated dimer (Figure 6D).^55^

The most well-characterized molecular mechanism of Orai activation is attributed to SOCE, which requires [Ca^2+^]_ER_ depletion and the interaction STIM. While SOCE has been shown to occur on the apical membrane of polarized epithelial cultures^53^, the biophysical parameters and the relative magnitudes of apical vs. basolateral Orai activation are unresolved. As a “proof of principle” experiment to determine if apical membrane activation of Orai can enhance ciliary beating, we tested the synthetic Orai1-specific agonist IA65 to stimulate ciliary beating.^56^ When treated with IA65 on the apical side only, human nasal ALI cultures showed an increase in the percent area of beating cilia compared to application of vehicle control, shown as an increase in the active area of beating cilia (Figure 6E,F,G, and Figure S11). Ciliary beat measurements demonstrated an increase in CBF, along with a recruitment of beating cilia (Figure 6H). We corroborated the activation of beating cilia as physiologically meaningful by a particle transport assay that approximates changes in mucociliary clearance.^57,58^ Fluorescent beads were added to the apical side of ALI cultures to measure the distance beating cilia could propel particles when challenged with IA65. Streak lengths over a fixed time interval were used to calculate particle velocity. HNEC ALIs showed a 30-50% increase in particle transport after IA65 (Figure 6I,J). These results demonstrate Orai1 is localized to airway motile cilia, and apically localized Orai1 can be targeted to modify cilia beating.

### Motile cilia are stimulated by arachidonic acid through a mechanism potentially involving Orai1 and MS4A8B

While canonical Orai activation occurs during [Ca^2+^]_ER_ depletion and STIM interaction with Orai, Orai channels are also activated in a store-independent manner by arachidonic acid (AA) generated from phospholipids; AA is thus thought to be a physiological ligand for Orai.^59,60^ Ciliary membranes are rich in phosphoinositol that can be a precursor to AA^61^, and AA metabolism occurs in airway cells^62,63^ as a key component of both inflammatory and anti-inflammatory signaling. AA homeostasis may be disrupted in airway diseases like cystic fibrosis^64^, aspirin-exacerbated respiratory disease^65^, and asthma.^66^ We hypothesized AA may be a physiological ligand to activate cilia-specific pools of Orai1 which may be regulated by MS4A8B. When AA was applied exogenously to ciliated ALI cultures, CBF increased and was reversed with Orai Ca^2+^ channel inhibitor 2-APB (Figure 7A,B,C). AA-stimulated CBF was further inhibited by Orai1 inhibitor GSK-7975A^67,68^ (Figure S12). We hypothesized the increase in CBF would increase mucociliary clearance rates. Longer fluorescent streaks after AA challenge again demonstrated increased particle velocity and greater mucociliary clearance (Figure 7D,E).

**Figure 7:**
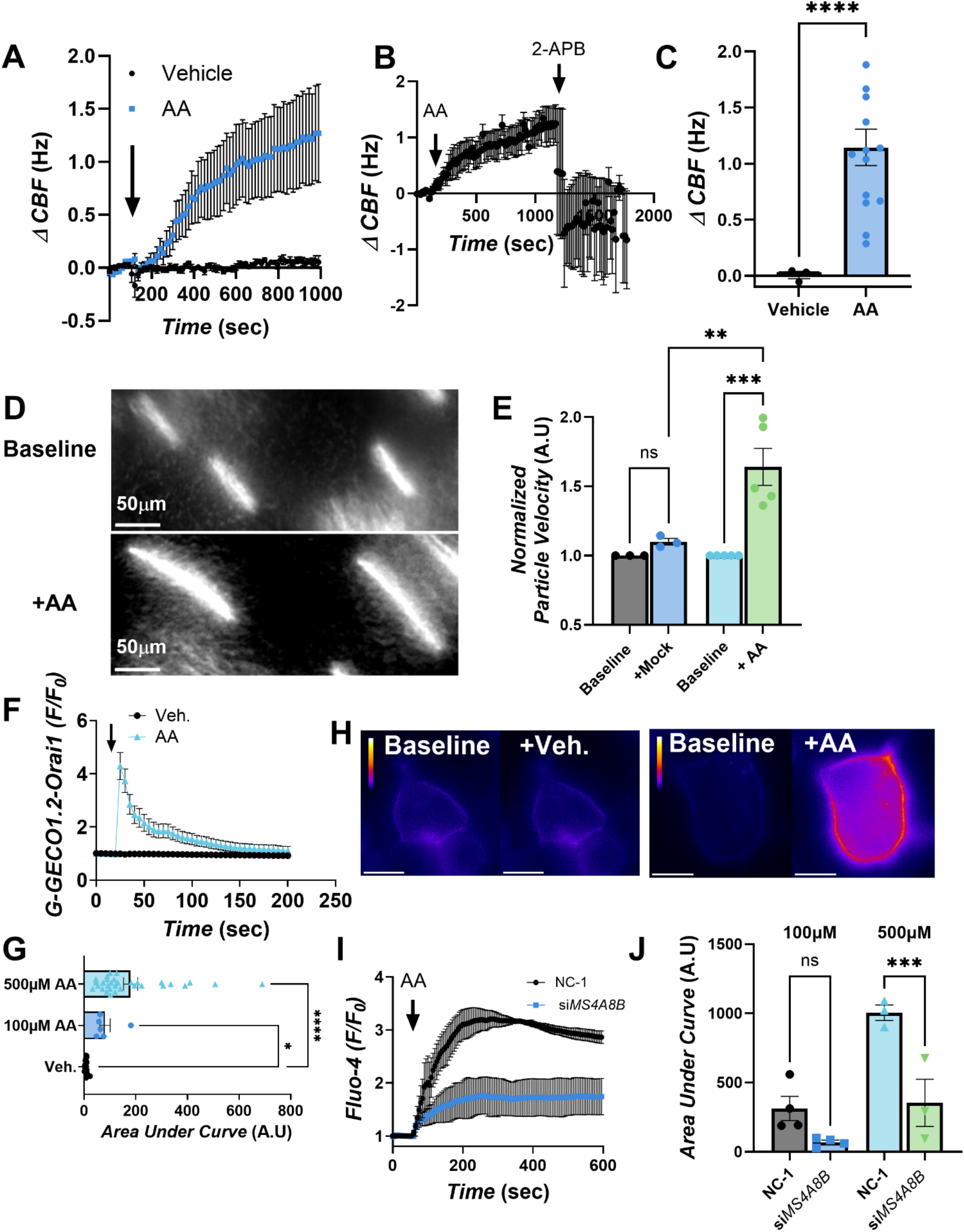
Arachidonic acid stimulates airway motile cilia dependent on Orai1. (A) CBF time course (mean ± SEM) of primary HNEC ALIs. Arachidonic acid (AA, 10μM). n=3 independent patients. (B) CBF time course (mean ± SEM) of primary HNEC ALIs stimulated with AA (10μM), and challenged with 2-APB (50μM). n=3. (C) Summary statistics of peak change in CBF after (AA, 10μM) or vehicle (<0.001% ethanol) taken within 8min of treatment (mean ± SEM). Vehicle, n=3; +AA n=14 independent patients. Significance by Welch’s unpaired t-test, p=<0.0001. (D) Particle transport micrographs from HNEC ALI cultures before and after AA (10μM). (E) Summary statistics of particle transport assay (mean ± SEM). Mock, n=3; +AA n=5 independent patients. Significance tested with a one-way ANOVA with Tukey’s multiple comparisons test, **p=0.0051, ***p=0.0004. (F) Time course of AA activated Ca^2+^ influx reported by G-GECO1.2-Orai1 in DMS53 cells (mean ± SEM). Vehicle (n=15 cells), +AA (500μM, n=31 cells). (G) Summary statistics of (F) indicating AUC (mean ± SEM) for each replicate. Significance by Brown-Forsythe and Welch ANOVA with Dunnett’s multiple comparison test, *p=0.0299; ****p=<0.0001. (H) Representative images of peak Ca^2+^ reported by G-GECO1.2-Orai1 after mock treatment or 500μM AA in DMS53 cells. Scale bar, 10μm (I) Time course of AA (500μM) activated Ca^2+^ influx reported by Fluo-4 (mean ± SEM) in DMS53 cells expressing MS4A8B (NC-1) or siRNA-mediated knockdown (si*MS4A8B*), n=3 independent transfections for both. (J) Summary statistics of Ca^2+^ responses between MS4A8B expressing DMS53 cells and MS4A8B knockdown (mean ± SEM). Significance by one-way ANOVA and Fishers Least Significant Difference test. ***p=0.0008, ns p=0.0671. See also Figures S12 and S13.

To further evaluate if MS4A8B can modulate AA-activated Orai Ca^2+^ fluxes, we applied AA to DMS53 cells and monitored [Ca^2+^]_cyt_. AA induced reproducible Ca^2+^ responses at 100μM and 500μM. Although this is higher than needed to activate the motile cilia of primary HNECs (Figure S13A,B), this could be due to differences in DMS53 cell membrane or Orai channel stoichiometry. Nonetheless, this AA-induced Ca^2+^ signaling depended on extracellular Ca^2+^ (Figure S13C,D). Orai1’s specific involvement in the AA responses was demonstrated by imaging of G-GECO1.2-Orai1 Ca^2+^ indicator (Figure 7F,H,G), which revealed AA stimulated Orai-mediated Ca^2+^ influx. The AA-induced [Ca^2+^]_cyt_ signals were abolished by si*MS4A8B* transfection, suggesting MS4A8B plays a regulatory role in facilitating AA-Orai signaling in airway cells (Figure 7I,J; Figure S13E).

## Discussion

Motile cilia beating is a key component to mucociliary clearance as the innate defense against inhaled pathogens. While ciliary beating’s dependence on Ca^2+^ signaling is well established, the molecular identity of Ca^2+^ transporters, especially those localized to the ciliary membrane, has remained incompletely understood. Here, we identify MS4A8B as a cilia-specific protein whose loss impairs ciliary function. Our data show MS4A8B physically and functionally couples with Orai1 to regulate multiple aspects of motile cilia function, including a novel arachidonic acid-Orai1 signaling pathway. These results provide molecular insight, expanding our understanding of how airway epithelial cells regulate ciliary beating and mucociliary clearance.

A major conclusion of our study is Orai1’s localization and activation on the ciliary membrane. Traditionally recognized for its role in SOCE, Orai1’s prominent apical localization to motile cilia suggests a highly compartmentalized, likely store-independent, signaling role. The apical sub-ciliary space of airway-ciliated epithelial cells is densely packed with mitochondria^40^ and cytoskeleton networks^69^ that would severely limit the fast translocation of ER-resident STIM to Orai for the rapid regulation needed for ciliary beating. Our data show cilia-localized Orai1 can directly modulate ciliary beating when activated by specific agonist IA65 and, importantly, AA. We hypothesize AA is a physiological ligand for this cilia-localized Orai. We speculate Orai1 is a biophysically suitable Ca^2+^ channel for cilia. The volumetric space of an individual cilium is approximately 10,000 fold smaller than the cell body.^70^ Orai1’s small conductance (<1pS) might provide enough Ca^2+^ to elevate cilia Ca^2+^ but minimize Ca^2+^ overload^71^ by limiting spillover of cilia Ca^2+^ into the mitochondria directly below. While Ca^2+^ is necessary for proper mitochondrial function, mitochondrial Ca^2+^ overload is a well-characterized activator of apoptosis.^72,73^ We speculate Orai regulation of Ca^2+^ signaling in cilia may allow compartmentalized ciliary signaling to remain compartmentalized.

AA is generated from phosphoinositol-rich ciliary membranes during inflammatory or mechanical stress. We demonstrate AA robustly increases CBF in a store-independent manner reliant on both Orai1 and MS4A8B, suggesting a specialized mechanotransduction or chemosensory pathway wherein MS4A8B facilitates Orai1 activation by localized lipid mediators. This pathway likely allows motile cilia to auto-regulate their CBF autonomously from global ER Ca^2+^ signaling, providing a rapid, localized response to airway irritation or pathogen-induced membrane stress. AA activation of Orai on cilia and regulation of ciliary beating may thus represent early inflammatory alarms in the airway epithelium. AA is produced upstream of inflammatory mediators, including prostaglandins and leukotrienes converted by cyclooxygenase and 5-Lipooxygenase pathways, respectively. Single-nucleotide polymorphisms on chromosome 11q, which harbors coding sequences for the entire MS4A family, may be asthma risk factors.^74^ Moreover, Orai1 is upregulated in asthma^51^, which has also been tied to dysregulated AA signaling.^66^

Orai1, and more generally SOCE, is the most prominent mechanism for Ca^2+^ influx in non-excitable cells, particularly immune cells. MS4A4A was previously shown to interact with Orai1 in both T cells and mast cells.^25,27^ Our results indicate functional regulation of Orai1 Ca^2+^ influx is conserved outside of immune cells. Unlike others who have speculated that MS4A proteins are ion channels^26,47^, our data strongly support that MS4A8B is not itself a Ca^2+^ channel but rather regulates Orai1. Collectively, we speculate MS4A interaction with Orai underpins other MS4A-associated diseases. Continued demonstration of MS4A regulation of Orai Ca^2+^ channel proteins in cell-type specific contexts would be valuable in neurons^34^ where MS4A proteins may regulate odor chemoreception, microglial cells where MS4A homologs have documented roles in Alzheimer’s disease progression^75-77^, and in B-cells and macrophages where there is a strong clinical link to MS4A homologs in rheumatoid arthritis.^33,36^ Orai isoforms’ ubiquitous expression suggests dysregulated Orai signaling may be the root cause of these MS4A-linked phenotypes.

## Resource availability

### Lead contact

Requests for further information and resources should be directed to and will be fulfilled by the lead contact, Robert J. Lee

### Materials availability

All unique reagents generated in this study are available from the Lead Contact

### Data and code availability

Numerical source data used to generate figures or other raw data are included as supplemental material. This paper does not report original code.

## Acknowledgments

We thank M. Victoria (University of Pennsylvania) for excellent technical assistance and helpful discussions. Work for this project with funded by NIH R01 HL178613 and HL181155 as well as the Stone Fund for Rhinology Research (University of Pennsylvania, Department of Otorhinolaryngology).

## Author contributions

Conceptualization, A.A.S and R.J.L; investigation, A.A.S, R.M; J.S.R; formal analysis, A.A.S, R.M; J.S.R, ZM; methodology, A.A.S, R.M, Y.I.K, K.R, Z.M; J.N.P, N.D.A; writing – original draft, A.A.S, R.J.L; writing – review & editing, A.A.S, R.Z, Y.I.K, K.R, Z.M, J.N.P, N.D.A, R.J.L; resources, Y.I.K., K.R., J.N.P., N.D.A.; funding acquisition, N.D.A., J.N.P., R.J.L.

## STAR ★ METHODS

### Key resources table

| REAGENT or RESOURCE | SOURCE | IDENTIFIER |
| --- | --- | --- |
| **Antibodies** |  |  |
| anti-MS4A8B (3E6) | Kerafast | Cat#: EUC001 |
| anti-MS4A8B | Sigma-Aldrich | Cat#: HPA007318;<br>RRID:AB_1854139 |
| anti-β-TUBULIN IV (EPR16775) | Abcam | Cat#: ab179504 |
| anti-ORAI1 (3F6H5) | Thermo Fisher Scientific<br>(Invitrogen) | Cat#: MA5-15776;<br>RRID:AB_11153035 |
| anti-MUC5AC (45M1) | Thermo Fisher Scientific(Life Technologies) | Cat#: MA5-12178;<br>RRID:AB_10978001 |
| anti-GFP | Abcam | Cat#: ab6556;<br>RRID:AB_305564 |
| anti-GLUT1 (SPM498) | Thermo Fisher Scientific | Cat#: MS-10637-P0<br>RRID:AB_721074 |
| anti-mouse IgG, Alexa Fluor 488 | Thermo Fisher Scientific<br>(Molecular Probes) | Cat#: A-21202;<br>RRID:AB_141607 |
| anti-rabbit IgG, Alexa Fluor 546 | Thermo Fisher Scientific<br>(Molecular Probes) | Cat#: A10040;<br>RRID:AB_2534016 |
| anti-rabbit IgG, HRP Conjugated | Santa Cruz | Cat# SC-2004;<br>RRID: AB_631746 |
| anti-mouse IgG, HRP Conjugated | Santa Cruz | Cat# SC-2005;<br>RRID: AB_631736 |
| **Chemicals, peptides, and recombinant proteins** |  |  |
| 2-APB | Tocris Bioscience | Cat#: 1224 |
| Alexa Fluor 647 phalloidin | Thermo Fisher Scientific<br>(Molecular Probes) | Cat#: A22287 |
| Adenosine triphosphate, magnesium salt | Sigma-Aldrich | Cat#: A9187 |
| Arachidonic acid | Cayman Chemical | Cat#: 90010.1 |
| BTP2 | Cayman Chemical | Cat#: 13246 |
| Carbachol | Tocris Bioscience | Cat#: 2810 |
| Thapsigargin | Cayman Chemical | Cat#: 10522 |
| Chromotek GFP-Trap | Proteintech | Cat#: gtma |
| Fluoroshield mounting media with DAPI | Abcam | Cat#: ab104139 |
| Fluo-4 AM | AAT Bioquest | Cat#: 20550 |
| FluoSpheres Carboxylate-Modified Microspheres | Thermo Fisher Scientific<br>(Invitrogen) | Cat#: F8826 |
| Gadolinium chloride | Thermo Fisher Scientific | Cat#: 11286-06 |
| GSK-7975A | Millipore-Sigma | Cat#: 5.34351.0001 |
| IA65 | Millipore-Sigma | Cat#: SML4116 |
| Triton-X | Sigma-Aldrich | Cat#: T6878 |
| Saponin | Fisher Scientific | Cat#: AC41923-1000 |
| Normal donkey serum (NDS) | Abcam | Cat#: ab166643 |
| Bovine serum albumin (BSA) | Sigma-Aldrich | Cat#: A7030 |
| Paraformaldehyde | Fisher Scientific | Cat#: 50-980-495 |
| TriZol | Thermo Fisher Scientific | Cat#: 15596026 |
| CellTak | Corning | Cat#: 354240 |
| **Critical commercial assays** |  |  |
| High-Capacity cDNA Reverse Transcription Kit | Thermo Fisher Scientific (Applied Biosystems) | Cat#: 4368814 |
| Direct-zol RNA Miniprep | Zymo research | Cat#: R2050 |
| Lipofectamine 3000 Transfection Reagent | Thermo Fisher Scientific (Invitrogen) | Cat#: L3000015 |
| Lipofectamine RNAiMAX Transfection Reagent | Thermo Fisher Scientific (Invitrogen) | Cat#: 3778075 |
| **Experimental models: Cell lines** |  |  |
| DMS53 | ATCC | CRL-2062 |
| DMS454 | Millipore-Sigma | Cat#: 95062832 |
| DMS153 | ATCC | CRL-2064 |
| BEAS-2B | ATCC | CRL-9609 |
| HEK293 | ATCC | CRL-1573 |
| HEK293 Orai Triple Knockout | Applied Biological Materials | Cat#: T6490 |
| Primary human nasal epithelial cells | Hospital of the University of Pennsylvania |  |
| **Oligonucleotides** |  |  |
| DsiRNA targeting MS4A8B | IDT | hs.Ri.MS4A8.13.1 |
| Non-targeting DsiRNA | IDT | Cat#: 51-01-14-04 |
| Custom Antisense oligonucleotide targeting MS4A8B | IDT | Cat#: 9462 |
| Custom scramble Antisense oligonucleotide | IDT | Cat#: 9462 |
| TaqMan Assay for MS4A8B | Thermo Fisher Scientific | Hs00230227_m1 |
| TaqMan Assay for UBC | Thermo Fisher Scientific | Hs00824723_m1 |
| **Recombinant DNA** |  |  |
| pcDNA:D3cpv | Roger Tsien Lab | Addgene #36323 |
| pcDNA:D1ER | Roger Tsien Lab | Addgene #36325 |
| pEGFP-C2:G-GECO1.2-Orai1 | Michael Cahalan Lab | Addgene #73562 |
| pcDNA3.1:mCherry-MS4A8B | In house |  |
| pcDNA3.1:mCherry | In house |  |
| pcDNA3.1:MS4A8B-mCherry | In house |  |
| pcDNA3.1:MS4A8B-CFP | In house |  |
| pEGFP-N1:Orai1-YFP | Anjana Rao Lab | Addgene #19756 |
| pRP:MS4A8B-IRES-mCherry | Vector Builder |  |
| pRP:MS4A8B | Vector Builder |  |
| pRP:aa2-83 beta galactosidase (empty vector, EV) | Vector Builder |  |
| pCMV6:MS4A8B-GFP | In house |  |
| pIRES2-EGFP:MS4A8B | In house |  |
| **Software and algorithms** |  |  |
| μmanager | Open Source | 81 |
| ImageJ/Fiji | Open Source | 82 |
| MetaMorph | Molecular Devices |  |
| MetaFluor | Molecular Devices |  |
| QuantStudio 5 | Applied Biosystems, Inc |  |
| Sisson Ammons Video Analysis (SAVA) | Ammons Engineering |  |
| PRISM | Graph Pad |  |
| Pulse | HEKA Elektronik |  |
| Igor Pro | WaveMetrics |  |
| CaSiAn | Moein *et al.*, | 83 |

### Methods

#### Cell culture

Primary Human Nasal Epithelial Cells (HNEC) were collected from cytological brushing performed during sinonasal surgery, as previously reported^84^, following institutional review board approval (#800614) in accordance with The University of Pennsylvania guidelines for use of residual clinical material, the U.S. Department of Health and Human Services code of federal regulation Title 45 CFR 46.116, and the Declaration of Helsinki. Written informed consent was received for all samples. Primary HNECs were expanded in T-25 culture flasks with complete PneumaCult-Ex Plus medium containing 1%(v/v) Penicillin/Streptomycin mix, 0.125 µg/mL amphotericin B, 10 µM Rock inhibitor Y27632 and 1 µM DMH-1 to 80% confluency (P0), and transferred to Transwell® (Corning) cell culture inserts (0.33 cm^2^ PTFE membrane, 0.4 µm pore size, transparent) under submerged conditions. When primary HNECs achieved confluency on transwells (P1), apical media was removed and cells were differentiated at Air-Liquid Interface (ALI) for 21 days and fed complete PneumaCult ALI medium (Stemcell, supplemented with 1% Penicillin/Streptomycin mix, 0.125µg/mL amphotericin B) basolaterally. HNEC’s were maintained at ALI in humidified incubators at 37°C and 5% CO_2_.

DMS53 (ATCC, CRL-2062), DMS454 (Millipore-Sigma, 95062832), and DMS153 (ATCC, CRL-2064) were cultured in Waymouth’s MB 752/1 medium (Gibco #11220-035) with 10% Fetal Bovine Serum (FBS) and 1% Penicillin/Streptomycin mix. BEAS-2B (ATCC, CRL-9609) cells were cultured in Ham’s F-12K (Kaighn’s) medium (Gibco # 21127-022) with 10% FBS and 1% Penicillin/Streptomycin mix. HEK293 (ATCC, CRL-1573) and HEK293 Orai Triple Knockout (Applied Biological Materials, T6490) cells were grown in DMEM (Corning: #10-013-CV) with 10% FBS and 1% Penicillin/Streptomycin mix. All cell lines were incubated at 37°C and 5% CO_2_ and passaged at 80% confluency.

#### cDNA, DsiRNA, and ASO Transfection

Cells were lifted from T-25 cell culture flasks with 0.05% Trypsin (Gibco, # 25300054) and plated at 40% confluency in glass bottom 8-well chamber slides for transfection with Lipofectamine 3000 (Invitrogen) for transient cDNA over-expression, or transfected with RNAiMAX (Invitrogen) for siRNA transfection.

In all transient over-expression experiments, a transfection master mix was made with Lipofectamine 3000, cDNA, p3000 reagent in 250 μL Opti-Mem (Gibco, #31985062). 4 μl of Lipofectamine 3000 was used with following cDNA amounts. For membrane proteins and empty vector (EV) controls (pcDNA3.1:mCherry-MS4A8B, pcDNA3.1:MS4A8B-mCherry, pRP:MS4A8B, pCMV6:MS4A8B-GFP, pIRES2-EGFP:MS4A8B, pcDNA3.1:Orai1, pcDNA3.1:EV, pRP:EV), 1 μg cDNA was used. 0.5 μg cDNA was used for Ca^2+^ indicators pcDNA:D3cpv (Addgene: Plasmid #36323), pcDNA:D1ER (Addgene: Plasmid #36325). 0.75 μg cDNA was transfected for dual Ca^2+^ indicators and membrane protein (G-GECO1.2-Orai1). For FRET imaging, 1.5 μg total cDNA was transfected in a 1:1 ratio of MS4A8B-CFP to Orai-YFP (0.75 μg MS4A8B-CFP and 0.75 μg Orai1-YFP). 2 μL P3000 reagent was added per 1 μg of cDNA following the manufacturers instructions. Master mix was added to plated cells at a 1:10 dilution. Cells were imaged 48hrs post-transfection.

For DsiRNA tranfection, master-mixes containing 9 μL of RNAiMAX was incubated with 100 nM DsiRNA (IDT) in 300 μL Opti-Mem was incubated and added to plated cells at a 1:10 dilution. mRNA knockdown was validated 72 hrs post-transfections and functional assays were also conducted 72 hrs post-transfection. MS4A8B targeting sequence was as follows:

+ 5’ rArUrGrUrGrArGrUrGrUrCrArUrCrUrArUrCrCrArArArCAT 3’,

- 5’ rArUrGrUrUrUrGrGrArUrArGrArUrGrArCrArCrUrCrArCrArUrUrG 3’.

Non-targeting DsiRNA (IDT, #51-01-14-04) was used as a negative control.

Antisense oligonucleotides (ASOs) were synthesized by IDT using the following sequences. To target MS4A8B: +G*+C*+A*A*G*C*A*T*G*A*A*T*T*C*G*A*T*G*A*C*T*+T*+C*+A. Scramble control was synthesized as: +A*+A*+T*C*G*T*A*C*G*A*G*T*C*A*G*T*A*C*G*+C*+T*+A. Prior to transfection, fully differentiated primary HNECs grown at ALI were washed 3-4x with PBS to wash residual mucus. 1 μM ASO was combined with 4 μL Lipofectamine and 2 μL P3000 reagent in 50 μL Opti-Mem per ALI culture. Transfection master mix was applied directly to the apical surface for 4-6 hrs before removal. Cultures were transfected every third day for 12 days.

#### Ca^2+^ imaging

For CaMeleon based imaging (D3cpv and D1ER), imaging was performed at room temperature on a Nikon Eclipse TE2000-U microscope under 60x lens (Nikon Plan APO, 1.4NA, Oil immersion, 0.21 WD) with a metal halide illumination system (Prior Scientific, Lumens 200), and μmanager software. CaMeleon probes were excited under CFP excitation wavelength (430/24 nm) and emission was collected at CFP (470/24 nm) and YFP emission wavelengths (535/30 nm) with Chroma Technology filter set (#89009-ET) housed in a Lamda 10-3 optical filter set changer (Sutter Instruments) controlled through μmanager. Illumination laser power was set to 10%. YFP and CFP imaging were recorded with 200 ms exposure time. Images were generated using a Retiga R6 monochrome CCD camera (Teledyne). Images were analyzed offline with ImageJ/FIJI.

For Fluo-4 AM based imaging, cells were loaded with 5 μM Fluo-4 AM (AAT Bioquest; #20551) in standard Hank’s Balanced Salt Solution (HBSS) for 1hr. Live-cell imaging was performed same Nikon Eclipse imaging system on a 20x objective (Nikon Plan Fluor, 0.5NA, 2.1 WD). Fluo-4 was excited with standard single wavelength FITC excitation (470/40nm) and emission (525/50nm) with Chroma Technology filter set (#49002-ET). Illumination laser power was set to 25%. Ca^2+^ oscillation data was analyzed in a semi-automated manner with CaSiAn software.^83^

For G-GECO1.2-Orai1 imaging, we used an Olympus IX-83 imaging system at 60x (Olympus Plan APO 1.42NA, Oil immersion, 0.15 WD), X-Cite 120LED Boost LED illumination source (Excelitas), FITC filters (Chroma Technologies), Orca Flash 4.0 sCMOS camera (Hamamatsu) and MetaFluor software (Molecular Devices). illumination power was set to 25%, with 75 ms or 100 ms exposure time.

Ca^2+^ imaging solutions consisted of 0.49 mM MgCl_2_ hexahydrate, 0.41 mM MgSO_4_ heptahydrate, 5.3 mM KCl, 0.44 mM KH₂PO₄, 140 mM NaCl, 0.33 mM Na₂HPO₄ heptahydrate, 10 mM HEPES, pH 7.2 (NaOH) and 1.5 mM Ca^2+^ unless otherwise stated as Ca^2+^ free indicating the substitution of Ca^2+^ for 1 mM EGTA.

#### Ciliary Beat Measurements

Ciliary beating was measured at room temperature using a Nikon Eclipse TS100 microscope (Nikon Plan Fluor ELWD 20x0.45 objective), Basler acA1300-200u m Camera (Ahrensburg, Germany) coupled with a Nikon 0.7x DXM Lens (Minato city, Tokyo, Japan) and quantified using Sisson-Ammons Video Analysis (SAVA) software. Cilia were imaged at 120 frames per second on transparent transwells under submerged conditions with 30μL 1.3 mM Ca^2+^ bathing solution with 0.49 mM MgCl_2_ hexahydrate, 0.41 mM MgSO_4_ heptahydrate, 5.3 mM KCl, 0.44 mM KH₂PO₄, 140 mM NaCl, 0.33 mM Na₂HPO₄ heptahydrate, 10 mM HEPES, pH 7.2 (NaOH) apically.

#### Immunofluorescence

Isolation of HNECs cells from ALI cultures was performed by gently detaching the cells off the insert membranes using cytology brushes in HBSS and centrifuged at 13,000xg for 1 min. Cell pellets were resuspended in 0.25% Trypsin-EDTA (Gibco) at 37°C for 5 min treatment. Trypsin was neutralized in PneumaCult-Ex Plus medium and removed by centrifugation at 125xg for 5min. Cells were resuspended in HBSS and plated on 3% (v/v) Cell-Tak (Corning) coated chamber slides. Cells were allowed to adhere for 30 min before being fixation with 4% paraformaldehyde for 30 min at room temperature. HNECs were permeabilized in blocking buffer (PBS, 1% BSA, 1% normal donkey serum, 0.2% saponin) with 0.3% Triton-X for 1.5 hrs at 4°C. Primary antibodies were incubated (1:100 in blocking buffer) overnight at 4°C. Secondary antibodies labeling (Alexa Fluor-labeled, anti-mouse, anti-rabbit, 1:1000 in blocking buffer) was performed for 2 h at 4°C. Cells were washed 3x with PBS between each step including fixation, permeabilization, primary staining, secondary staining. Specific immunofluorescence signal was confirmed by incubation of secondary antibody-only control samples.

Cells were imaged in Fluoroshield Mounting Media (Abcam) on a Nikon IX-83 DSU spinning disk confocal under 60x (Olympus Plan APO 1.42NA, Oil immersion, 0.15 WD) with FITC and TRITC filters (Chroma Technologies), Orca Flash 4.0 sCMOS camera (Hamamatsu) and MetaMorph software (Molecular Devices). Images were analyzed in ImageJ.

For Immunofluorescence of intact ALI monolayers, fixation permeabilization, and antibody incubation was carried out on the apical side of the transwell keeping the basolateral side hydrated in PBS. Insert membranes were cut and mounted in Fluoroshield Mounting Media (Abcam) between microscopy slide and coverslip prior to imaging.

Primary antibodies used for immunofluorescence were as follows: β-Tub IV (Abcam, ab179504), MS4A8B (Kerafast, EUC001), MS4A8B (Sigma, HPA007318), Orai1 (Invitrogen, MA5-15776), MUC5AC (Life Technologies, MA5-12178).

#### Co-Immunoprecipitation and Western Blot

Co-immunoprecipitation was performed using magnetic ChromoTek GFP-Trap agarose beads conjugated to anti-GFP nanobodies (Proteintech). Cells were washed 3x in Ca^2+^-free (1mM EGTA) standard bathing solution and lysed using lysis buffer (10 mM Tris pH 7.5, 150 mM NaCl, 10 mM KCl, 0.5% deoxycholate, 0.5% Tween, 0.5% IGePawl, 0.1% SDS) and bath sonication. GFP-Trap slurry was equilibrated in Wash buffer (24.8 mM Tris-HCl, 24.8 mM Tris-Base, 150 mM NaCl, pH 7.5) and combined with clarified cell lysate material. Lysate and bead mixtures were rotated end-over-end at 4°C for 1 hr. Agarose beads were separated from solution with a magnetic block and washed 3x with Wash buffer. Washed GFP-nanobody conjugated beads were transferred a clean Eppendorf tube and resuspended in 2xSDS buffer (100 mM Tris/Cl pH 6.8, 20 % glycerol, 4% SDS, 0.04 % bromophenol blue, 10 % β-mercaptoethanol). Bound protein was eluted by heating at 98°C for 5min. Samples were run on a NuPage 4-12% Bis-TRIS gel (Invitrogen) with MES running buffer (Tris Base 50 mM pH 7.3, MES 50 mM, SDS 0.1%, EDTA 1 mM).

Gels were transferred using Bis-Tris transfer buffer (25 mM bicine, 25 mM bis-tris, 1 mM EDTA, 10% methanol) to a nitrocellulose membrane (0.45 µM, Bio-Rad). Blots were blocked for 1 hr at room temperature or overnight at 4°C in TBST (24.8 mM tris acid & base, 150 mM NaCl, 0.5% Tween 20) with 5% milk. Primary antibodies were used at 1:1000 in TBST with 5% BSA overnight. HRP-conjugated chemiluminescent secondary antibodies were used at 1:10,000 in 5% milk for 1 hr. Blots were imaged using SuperSignal^TM^ West Pico PLUS Chemiluminescent Substrate (Thermo Scientific) using ChemDoc MP Imaging System (BioRad).

Primary antibodies for Western blotting were as follows: β-Tub IV (Abcam, ab179504), Orai1 (Sigma, O8254), GFP (Abcam, ab6556).

#### Cilia Harvesting

Motile cilia from primary HNECs were harvested from mature ALI cultures by detergent agitation as previously reported.^54^ Briefly, stock buffer (10mM Tris-HCl, 50 mM NaCl, 10 mM CaCl_2_, 1 mM EDTA, pH7.5) was used to create a deciliation buffer by spiking with 0.1% Triton X-100, 7mM β-mercaptoethanol, and 1% protease inhibitor cocktail (Roche) immediately before use. The apical side of ALI cultures were submerged in 300μL deciliation buffer and manually rocked for 1min before collecting detached cilia. Rocking with fresh deciliation buffer was repeated for a total of 3 rounds. Cilia were separated from cell debris by slow centrifugation at 1000xg for 1min. Cilia containing supernatant was transferred to clean Eppendorf tube and collected by centrifugation at 12,000xg. Cilia pellets were lysed using standard lysis buffer lysis buffer (10 mM Tris pH 7.5, 150 mM NaCl, 10 mM KCl, 0.5% deoxycholate, 0.5% Tween, 0.5% IGePawl, 0.1% SDS) and bath sonication.

#### FRET Imaging

FRET imaging was performed using an Olympus IX-83 widefield microscope at 60x (Olympus Plan APO 1.42NA, Oil immersion, 0.15 WD), on BEAS-2B cells transiently transfected with MS4A8B-CFP and Orai1-YFP, utilizing the acceptor photobleaching method. At each time point prior to bleaching, both the CFP channel (excitation 436/20 nm with 455 long pass dichroic, emission 470/24 nm) and the FRET channel (CFP excitation 436/24 nm, YFP emission 535/30 nm) were recorded. Subsequently, YFP photobleaching was conducted using excitation at 500/20 nm combined with a 515 nm long-pass dichroic filter and 535/30 emission filter, utilizing 10-second exposures at 100% LED power.

FRET efficiency (E_eff_) was calculated with the following equation according to^85^:

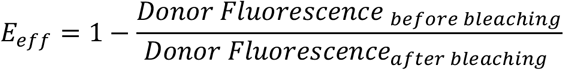

Proximity between donor and acceptor (R) was calculated by:

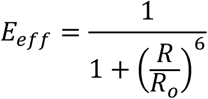

where R_o_=4.9nm^48^

#### RNA isolation and Quantitative reverse transcription PCR (qPCR)

RNA isolation was performed with the Direct-zol^TM^ RNA kit (Zymo Research) according to the manufacturer’s instructions using TRIzol^TM^ for homogenization. Complementary DNA (cDNA) was synthetized using the High-Capacity cDNA Reverse Transcription Kit (ThermoFisher Scientific) and quantified using TaqMan qPCR probes (ThermoFisher Scientific) in a QuantStudio 5 Real-Time PCR System.

#### Particle Transport Assay

Particle transport assay was completed as previously described.^57,58^ 0.2μm carboxylate-modified FluoSpheres (Invitrogen; F8826) diluted to 1:10,000 in HBSS were added to apical region on HNEC ALI cultures. Imaging was performed on a widefield fluorescent Nikon Eclipse TE2000-U microscope using TRITC excitation and excitation using 2.0 sec exposure times under 10x (Nikon Plan Fluor, 0.3NA, 16.0 WD). Transport velocity was normalized based on measurements within discrete regions of interest. Measurements were taken at baseline prior to treatment and 5-10 min after.

