## Supplemental Material for "MS4A8B regulates Orai1-dependent Ca²⁺ influx to control motile cilia function in human nasal epithelial cells"

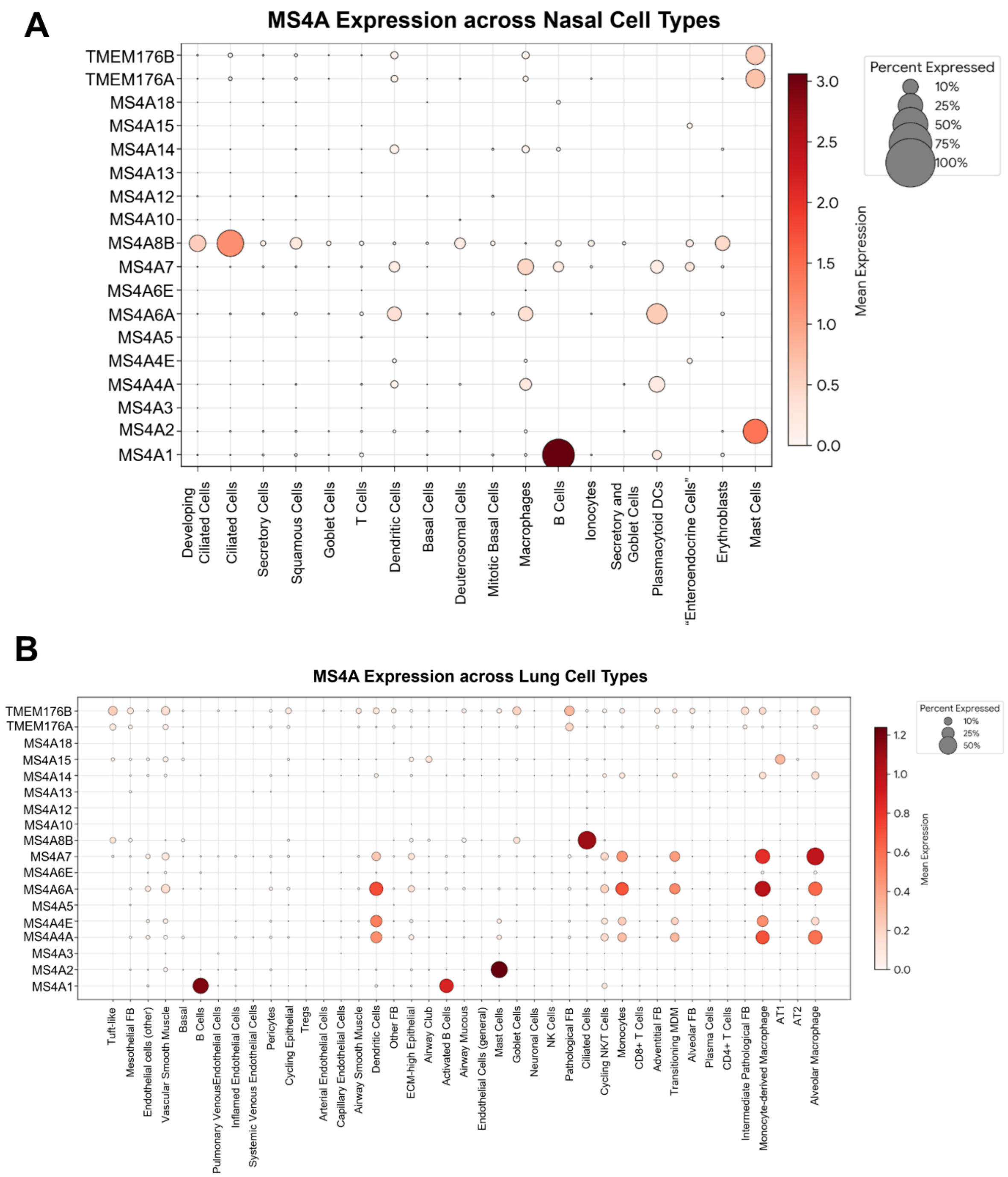


**S1: MS4A gene expression in nasal and lung tissue. Related to Figure 1.**

(A) Single-cell RNA sequence data highlighting MS4A homolog expression in nasal tissue. Data were accessed via Broad Institute Single Cell Portal and Ziegler et al.,[s1,s2]

(B) Single-cell RNA sequence data highlighting MS4A homolog expression in lung tissue. Data were accessed via Broad Institute Single Cell Portal and Melms et al.[s1,s3]

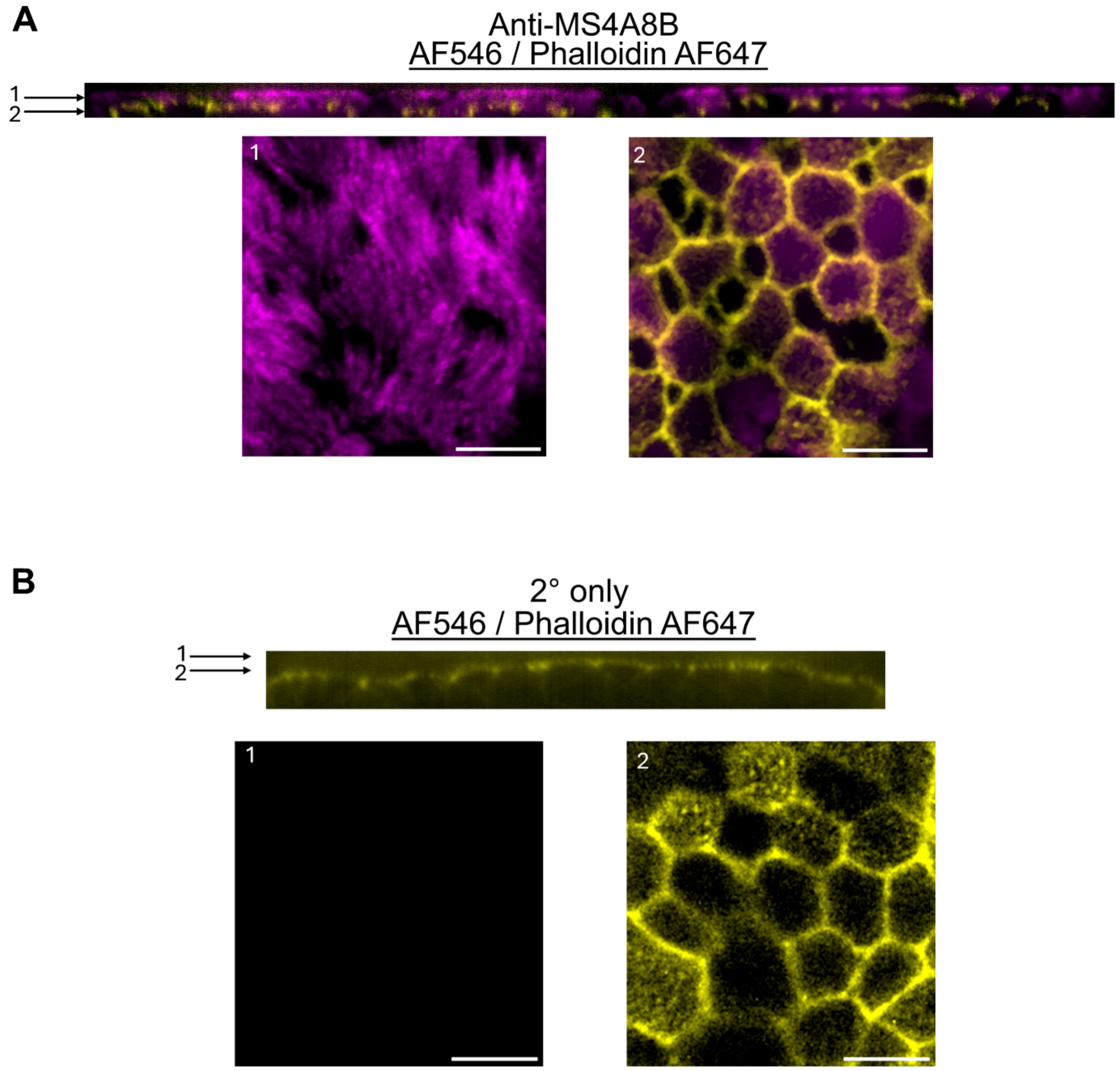


**S2: MS4A8B localization to motile cilia in primary HNECs of intact ALI cultures. Related to Figure 1.**

(A) Representative immunofluorescent image of MS4A8B on intact ALI culture. (Top) Orthogonal view of ALI. Bottom panel 1: Visualization of apical region of ALI culture, Bottom panel 2: Visualization of basolateral portion of ALI. Anti-MS4A8B conjugated to Alexa Fluor 546 (AF546) pseudo-colored with magenta, and phalloidin conjugated to Alexa Fluor 647 (AF647) pseudo-colored in yellow. Scale bar denotes 10μM.

(B) Immunofluorescence of HNEC ALI cultures labeled only with secondary antibodies. Showing an orthogonal view as done in (A) and visualizations of apical regions (1) and basolateral region (2), also as done in (A). Scale bar denotes 10μM.

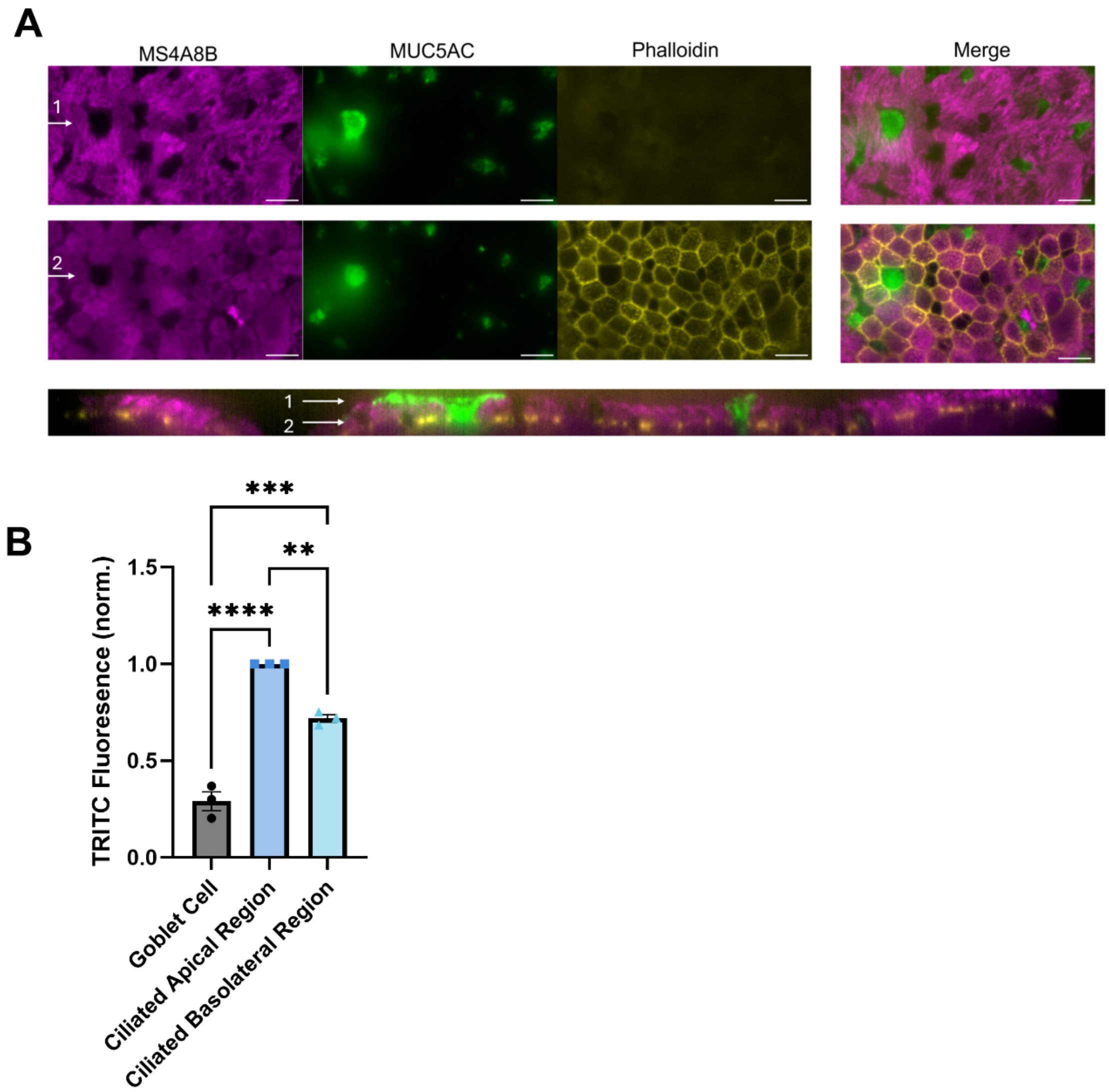


**S3: MS4A8B localization on HNEC ALI cultures. Related to Figure 1.**

(A) Immunofluorescence of MS4A8B compared to goblet cell marker MUC5AC, and F-actin marker phalloidin on the apical side of ALI cultures (1) and basolateral side (2). Bottom panel shows orthogonal view corresponding to apical and basolateral visualizations. Scale bars denote 10$\mu$m.

(B) Quantification of TRITC fluorescence exciting secondary Alexa Fluor 546 antibodies conjugated to MS4A8B. Error bars show S.E.M. Significance test with one-way ANOVA with Tukey's multiple comparisons test. ****p=<0.0001, ***p=0.0001, **p=0.0014.

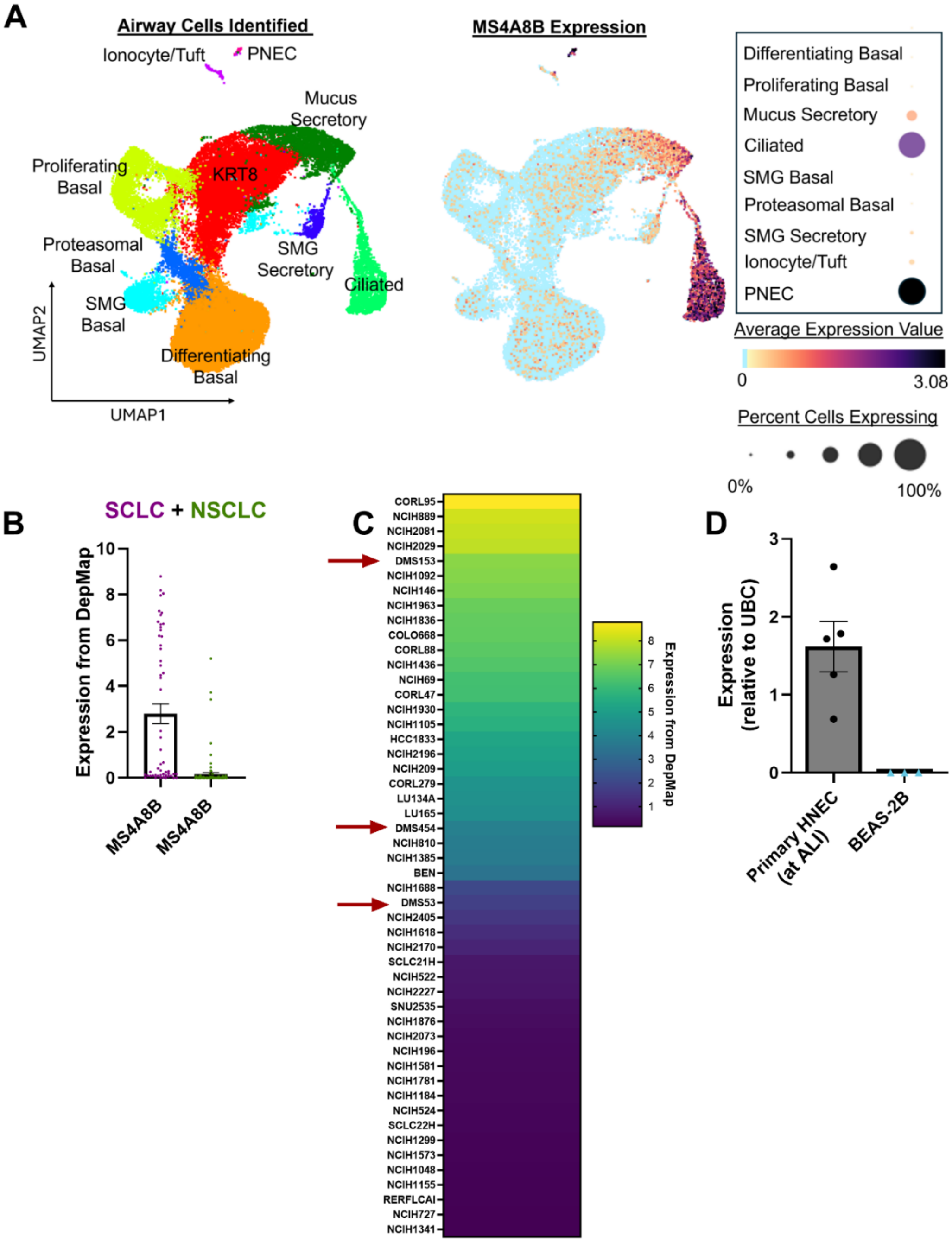


**S4: MS4A8B is expressed in tracheal ciliated cells and pulmonary neuroendocrine cells (PNECs), and PNEC-derived small cell lung cancer cells. Related to Figure 3.**

(A) UMAP projection of tracheal cell types identified with their corresponding MS4A8B expression intensity (36,248 total cells analyzed). Data were accessed from UCSC Cell Browser and Goldfarbmuren et al,.[s4,s5]

(B) Comparison of MS4A8B expression in small cell lung cancer (SCLC, magenta) and non-small cell lung cancer (NSCLC, green) accessed from DepMap.[6,7] Each data point represents an individual cell line. Error bars indicate S.E.M.
(C) Heat map illustrating the 50 most highly expressing MS4A8B cell lines derived from lung cancer cell lines found in DepMap. Red arrows point to DMS53, DMS454, and DMS153 selected as representative cell lines for subsequent functional assays.
(D) qPCR of MS4A8B transcripts in primary HNECs grown at ALI compared to submerged BEAS-2B cells. Error bars indicate S.E.M. For HNECs, expression was tested from 5 unique patients (n=5). For BEAS-2Bs, expression was tested from 3 separate passages (n=3).

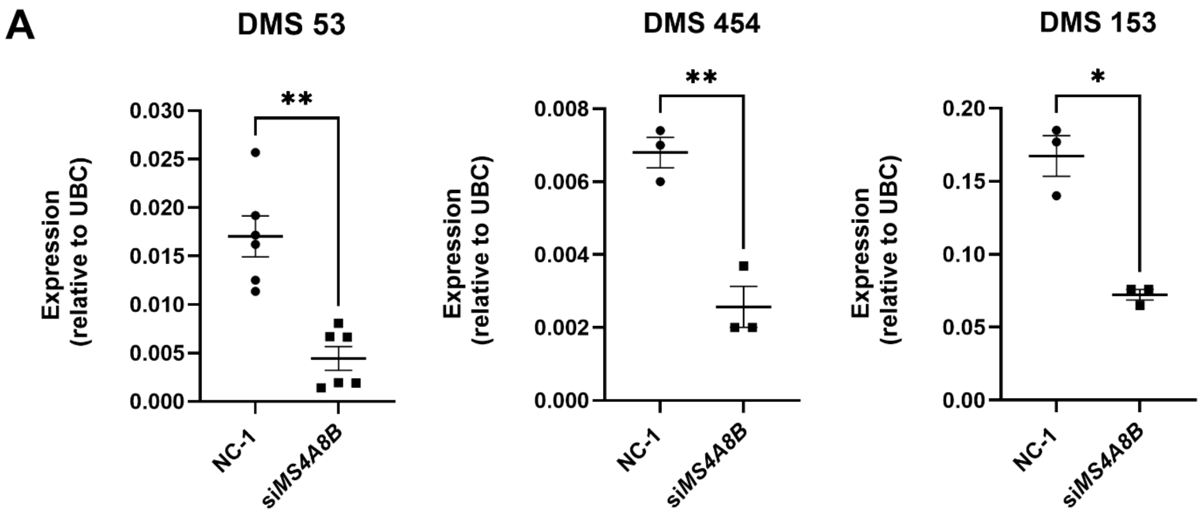


**S5: siRNA reduces MS4A8B mRNA expression in three independent airway epithelial cell lines. Related to Figure 3.**

Quantification of MS4A8B mRNA expression by qPCR following siRNA transfection with negative control (NC-1) or MS4A8B (si*MS4A8B*) targeting siRNA. Each point represents biological replicate, n=3-6. Error bars indicate S.E.M. Statistical significance tested with a paired t-test. For DMS53, **p=0.0047; for DMS 454, **p=0.0083; for DMS 153, *p=0.0287.

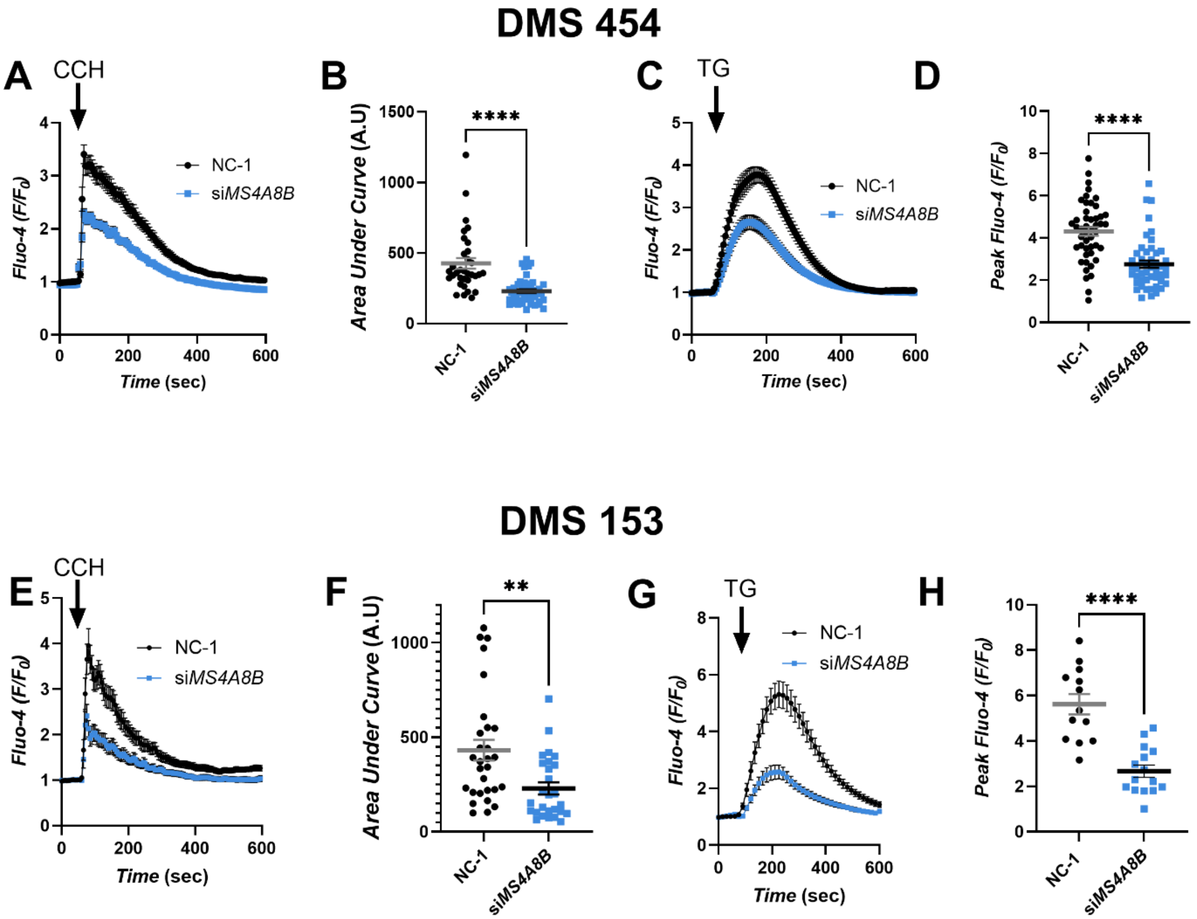


**S6: MS4A8B regulates ER $Ca^{2+}$ in DMS 454 and DMS 153 cells. Related to Figure 3.**

(A) Time course of $[Ca^{2+}]_{cyt}$ indicated by Fluo-4 signal in DMS 454 cells transfected with non-coding siRNA (NC-1, black line, n=33) or si*MS4A8B* (blue line, n=45) treated with carbachol (CCH) (10μM). Error bars represent S.E.M.

(B) Summary statistics corresponding to (A) showing the area under the curve (AUC) of Fluo-4 reported $Ca^{2+}$ traces of individual cells. Error bars represent S.E.M. Statistical significance tested with Welch's unpaired t-test, ****p= <0.0001.

(C) Time course of $[Ca^{2+}]_{cyt}$ indicated by Fluo-4 signal in DMS 454 cells transfected with non-coding siRNA (NC-1, black line, n=47) or si*MS4A8B* (blue line, n=47) treated with thapsigargin (TG) (10μg/mL). Cells were imaged in $Ca^{2+}$-free bath solution containing 1mM EGTA. Error bars represent S.E.M.

(D) Summary statistics of (C) showing the peak Fluo-4 signal. Error bars represent S.E.M. Statistical significance tested with Welch's unpaired t-test, ****p= <0.0001.

(E) Time course of $[Ca^{2+}]_{cyt}$ indicated by Fluo-4 signal in DMS 153 cells transfected with non-coding siRNA (NC-1, black line, n=29) or si*MS4A8B* (blue line, n=27) treated with carbachol (CCH) (10μM). Error bars represent S.E.M.

(F) Summary statistics corresponding to (E) showing the area under the curve (AUC) from Fluo-4 reported $[Ca^{2+}]_{cyt}$ traces of individual cells. Error bars represent S.E.M. Statistical significance tested with Welch's unpaired t-test, **p= <0.0047.

(G) Time course of $[Ca^{2+}]_{cyt}$ indicated by Fluo-4 signal in DMS 153 cells transfected with non-coding siRNA (NC-1, black line, n=13) or si*MS4A8B* (blue line, n=22) treated with thapsigargin (TG) (10μg/mL). Cells were imaged in $Ca^{2+}$-free bath solution containing 1mM EGTA. Error bars represent S.E.M.

(H) Summary statistics of (G) showing the peak Fluo-4 signal. Statistical significance tested with Welch's unpaired t-test, ****p= <0.0001.

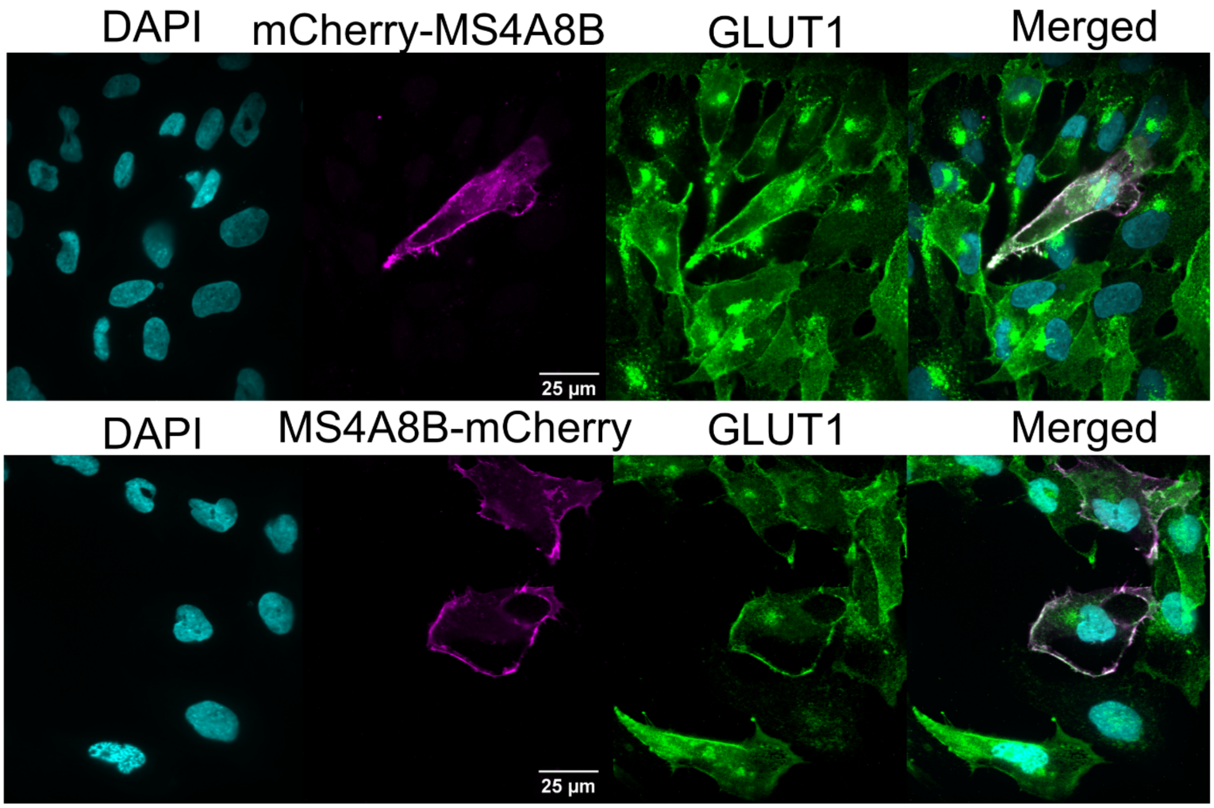


**S7: mCherry tagged MS4A8B constructs localize to the plasma membrane in BEAS-2B cells. Related to Figure 4.**

Representative immunofluorescence images comparing MS4A8B localization to plasma membrane marker GLUT1.

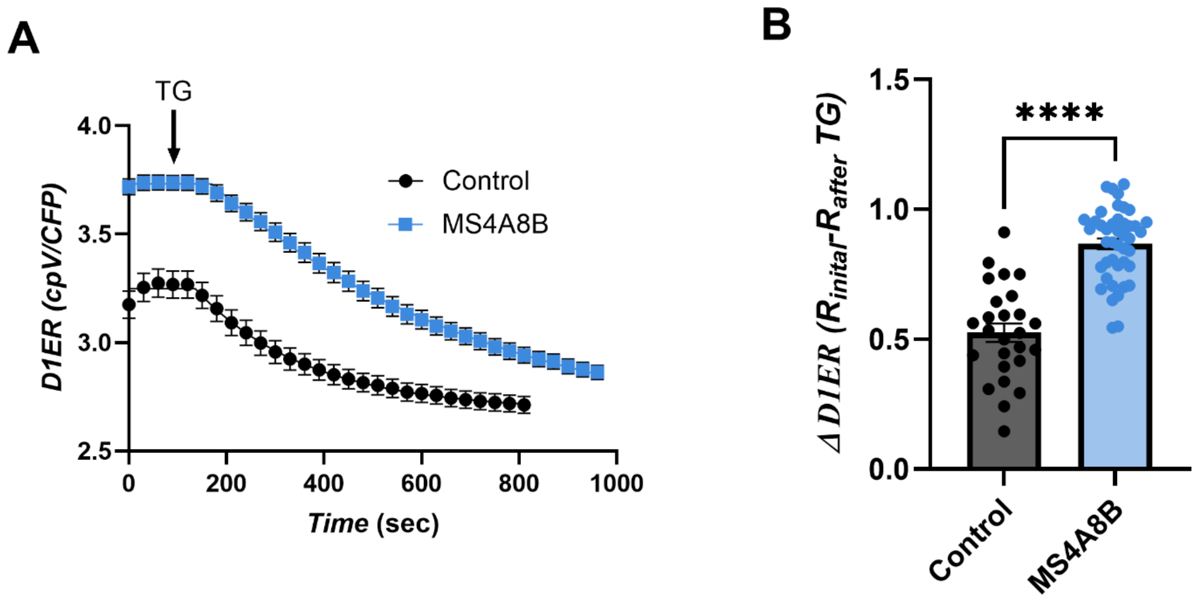


**S8: MS4A8B expression in HEK293 cells increases $[Ca^{2+}]_{ER}$. Related to Figure 4.**

(A) Average time course of $[Ca^{2+}]_{ER}$ reported by D1ER at rest and with stimulation by thapsigargin (TG, 10µg/mL). Control (black line, n=26), MS4A8B (blue line, n=43). Error bars indicate S.E.M. Cells were imaged 72hrs after transfection with D1ER and empty pRP vector, or D1ER + pRP:MS4A8B.

(B) Summary statistics showing the total change in D1ER fluorescence for each cell. Error bars indicate S.E.M. Statistical significance tested using unpaired t-test, ****p=<0.0001.

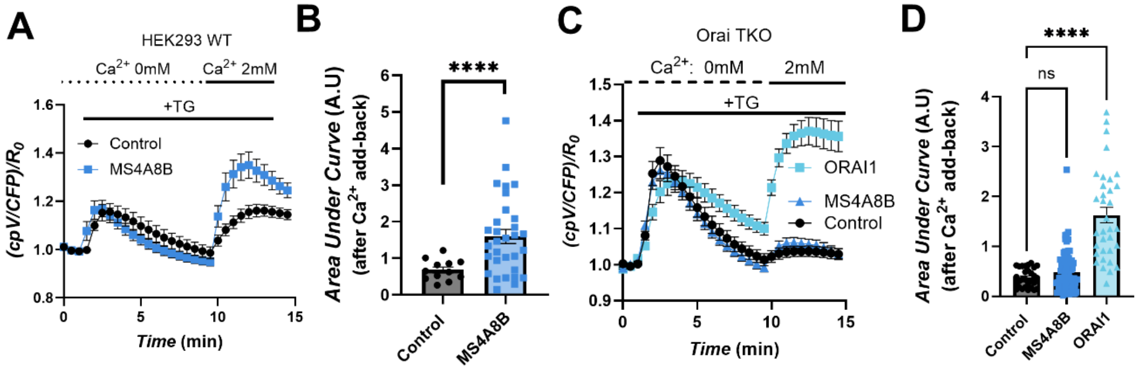


**S9: MS4A8B potentiation of store-operated $Ca^{2+}$ entry requires Orai homologs. Related to Figure 4.**

(A) Average time course of $[Ca^{2+}]_{cyt}$ reported by D3cpv in HEK293 cells transfected with empty vector control (black line, n=12) or MS4A8B (blue line, n=31) following ER $Ca^{2+}$ store depletion with 1mM EGTA plus 10μg/mL TG and $Ca^{2+}$ addback. Error bars show S.E.M

(B) Summary statistics analyzing area under curve of each cell imaged. Error bars show S.E.M. Statistical significance tested with Welch's unpaired t-test, ****p=>0.0001.

(C) Average time course of $[Ca^{2+}]_{cyt}$ reported by D3cpv in Orai triple knockout (TKO) HEK293 cells transfected with empty vector control (black line, n=29) or MS4A8B (blue line, n=80) or Orai 1 (cyan line, n=34) following ER $Ca^{2+}$ store depletion with 1mM EGTA plus 10μg/mL TG and $Ca^{2+}$ addback. Error bars show S.E.M

(D) Summary statistics analyzing area under curve of each cell imaged. Error bars show S.E.M. Statistical significance tested with one-way ANOVA with Dunnett's multiple comparison test, ****p=<0.0001.

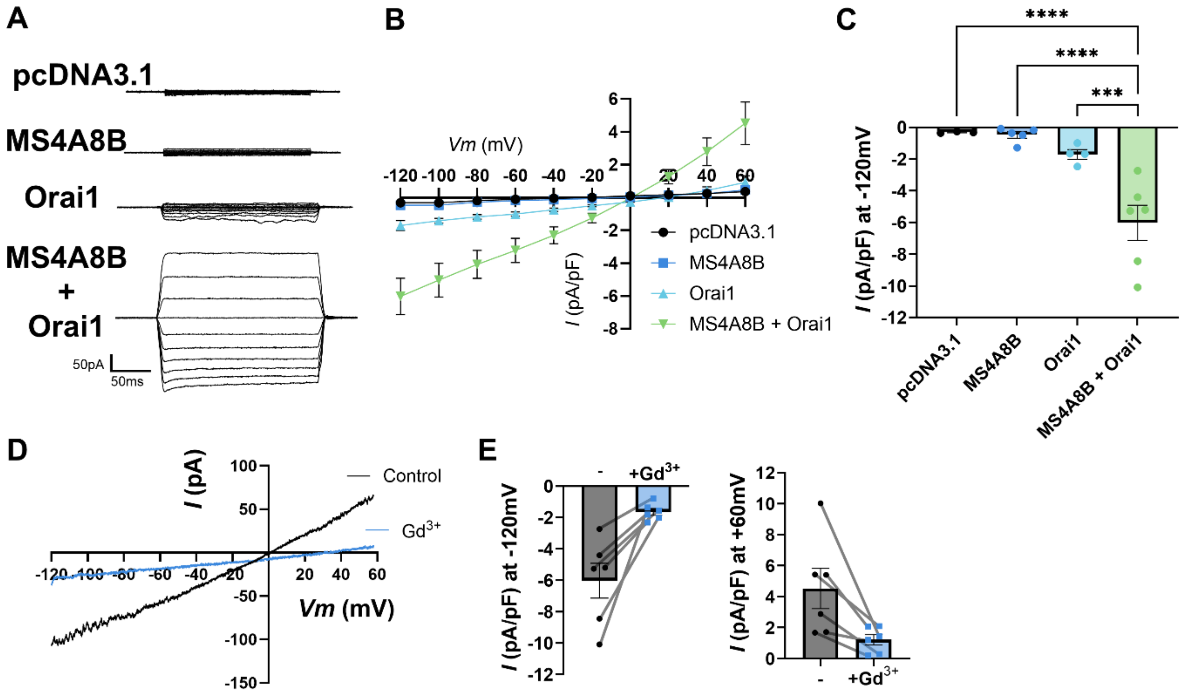


**S10: Orai1 is required for MS4A8B potentiated $Ca^{2+}$ current. Related to Figure 4.**

(A) Representative whole-cell currents for pcDNA3.1 vector (control), MS4A8B-GFP, Orai1-YFP, and co-expression of Orai1-YFP with MS4A8B (Orai1-YFP + MS4A8B) in Orai1/2/3 triple-knockout HEK293 cells.

(B) *I-V* relations for control (pcDNA3.1 vector, black circles, n=3, pF=22.67± 4.3), MS4A8B-GFP (blue squares, n=5, pF=28.0± 2.0) Orai1-YFP (cyan triangles, n=4, pF=28.0± 2.3, and co-expressed Orai1-YFP + MS4A8B (green triangles, n=6, pF=24.5±2.2). Error bars indicate S.E.M.

(C) Normalized currents (pA/pF) at -120 mV for individual cells. Error bars indicated S.E.M. Significance test by one-way ANOVA with Dunnett's multiple comparison, ****p=<0.0001, ***p=0.003.

(D) Whole-cell current traces from Orai TKO cell co-expressing MS4A8B and Orai1 evoked by a voltage-ramp protocol in absence (black line) and presence of 100 μM $Gd^{3+}$ (blue line).

(E) Individual cell changes in current amplitude given 100 μM $Gd^{3+}$ treatment (n=6). Error bars indicate S.E.M.

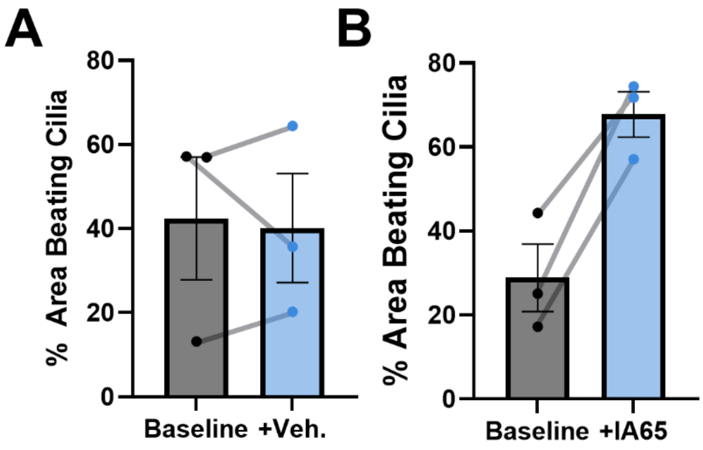


**S11: Patient-matched changes in ciliary beating after IA65 treatment. Related to Figure 6.**

Changes in area of beating cilia defined on an individual patient basis after vehicle treatment (0.05% DMSO) or IA65 (1μM). Error bars represent S.E.M.

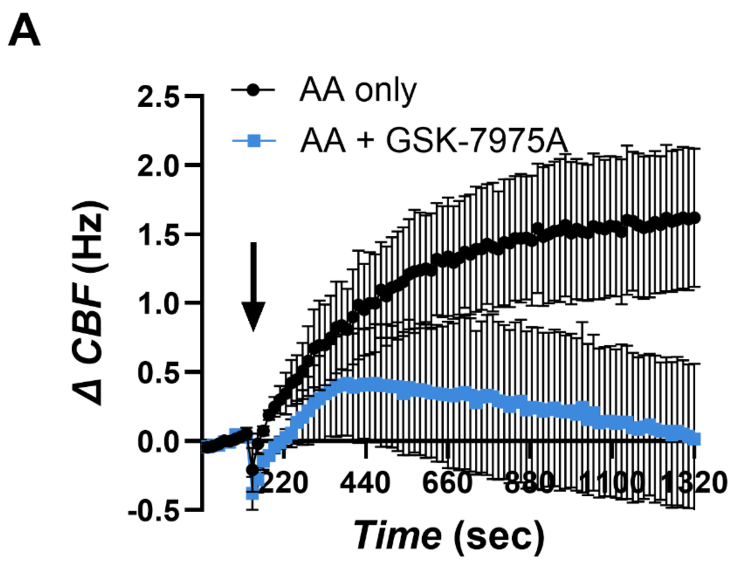


**S12: Arachidonic acid (AA) responses are inhibited by GSK-7975A. Related to Figure 7.**

(A) Average time course of AA induced changes to ciliary beat frequency (CBF) when treated with AA alone (10μM black, n=4) or co-treated with GSK-7975A (100μM; blue, n=4). Error bars represent S.E.M.

(B) Individual patient-matched differences in CBF corresponding to (A). *p=0.04 determined by paired t-test.

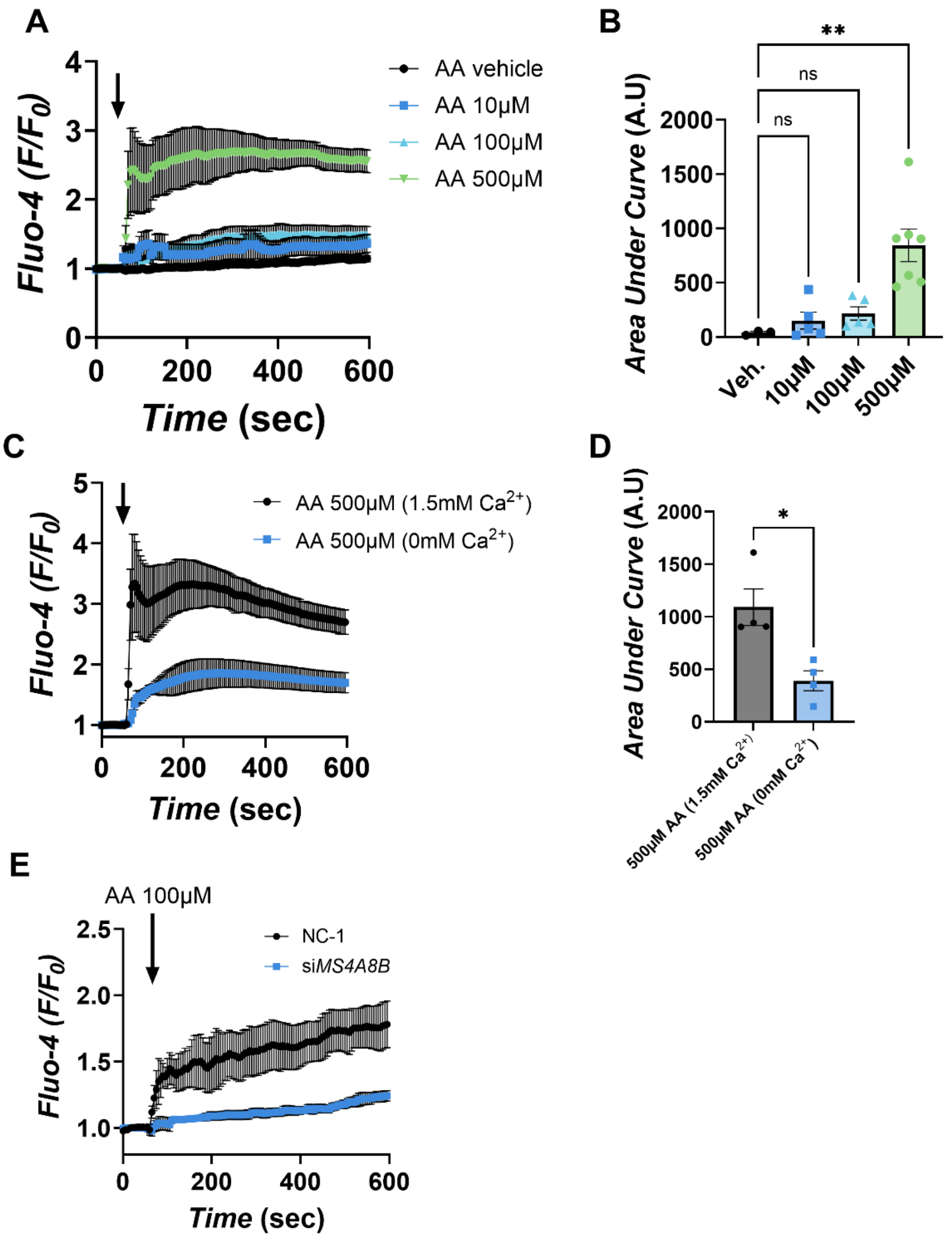


**S13: Arachidonic acid (AA) stimulation of DMS 53 cells. Related to Figure 7.**

(C) Average time course of $[Ca^{2+}]_{cyt}$ reported by Fluo-4 in DMS 53 cells when stimulated with AA at 10µM (blue squares, n=5), 100µM (cyan triangles, n=5), 500µM (green triangles, n=7), or vehicle control (black circles, n=3). Each "n" represents an independent transfection. Error bars represent S.E.M.

(D) Summary statistics detailing area under the curve for each sample. Error bars represent S.E.M. Significance tested with Brown-Forsythe and Welch ANOVA with Dunnett's multiple comparison test, **p=0.0048.

(E) Average time course of $[Ca^{2+}]_{cyt}$ reported by Fluo-4 in DMS 53 cells stimulated by 500μM arachidonic acid comparing bathing solution with 1.5mM $Ca^{2+}$ (black line, n=4 independent transfections) and $Ca^{2+}$ free (1mM EGTA) solution (blue line, n=4 independent transfections). Error bars represent S.E.M.

(F) Summary statistics detailing $Ca^{2+}$ responses from individual replicates. Error bars represent S.E.M. Significance test by Welch's unpaired t-test, *p=0.0188.

(G) Average time course of 100μM arachidonic acid evoked $[Ca^{2+}]_{cyt}$ reported by Fluo-4 in DMS 53 cells expressing MS4A8B having been transfected with negative control siRNA (NC-1, black line, n=4 independent transfections), or under gene siRNA mediated gene knockdown (si*MS4A8B*, blue line, n=4 independent transfections). Error bars represent S.E.M.

### Supplementary Methods

#### Electrophysiology

Whole cell currents in Orai triple knockout HEK293 cells were recorded 24 hours after transfection with MS4A8B-GFP, Orai-YFP, both constructs, or empty vector. Data was acquired with an Axopatch 200B amplifier (Axon Instruments) with an ITC-16 interface (Instrutech). Currents were filtered by low-pass Bessel filter at 1 kHz and sampled at 5 kHz. Electrode capacitance was compensated electronically. HEKA Pulse software (HEKA Elektronik) was used for data acquisition and stimulation protocols. Igor Pro was used for graphing and data analysis (WaveMetrics). Pipettes were pulled with a P97 puller (Sutter Instrument) to a resistance of 3 – 5 MΩ. Bathing solution contained 125 mM NaCl, 20 mM $CaCl_2$, 1 mM $MgCl_2$, 10 mM Glucose, 10 mM HEPES, pH 7.4 adjusted by CsOH. Pipette solution contained 120 mM CsCl, 8 mM $MgCl_2$, 11 mM EGTA, 10 mM HEPES, pH 7.3 adjusted by methanesulfonic acid. Following establishment of whole-cell configuration, internal solution was dialyzed for 7-10 min for the stable steady-state whole-cell currents. The whole-cell currents were evoked by 200-ms voltage-pulses from -120 mV to +60 mV in +20 mV increments from a holding potential of 0 mV. Voltage-ramp protocol was also run from -120 mV to +60 mV over 2 seconds. Currents were normalized to individual whole-cell capacitance.
